# Improving short and long term genetic gain by accounting for within family variance in optimal cross selection

**DOI:** 10.1101/634303

**Authors:** Antoine Allier, Christina Lehermeier, Alain Charcosset, Laurence Moreau, Simon Teyssèdre

## Abstract

The implementation of genomic selection in recurrent breeding programs raised several concerns, especially that a higher inbreeding rate could compromise the long term genetic gain. An optimized mating strategy that maximizes the performance in progeny and maintains diversity for long term genetic gain on current and yet unknown future targets is essential. The optimal cross selection approach aims at identifying the optimal set of crosses maximizing the expected genetic value in the progeny under a constraint on diversity in the progeny. Usually, optimal cross selection does not account for within family selection, i.e. the fact that only a selected fraction of each family serves as candidate parents of the next generation. In this study, we consider within family variance accounting for linkage disequilibrium between quantitative trait loci to predict the expected mean performance and the expected genetic diversity in the selected progeny of a set of crosses. These predictions rely on the method called usefulness criterion parental contribution (UCPC). We compared UCPC based optimal cross selection and optimal cross selection in a long term simulated recurrent genomic selection breeding program considering overlapping generations. UCPC based optimal cross selection proved to be more efficient to convert the genetic diversity into short and long term genetic gains than optimal cross selection. We also showed that using the UCPC based optimal cross selection, the long term genetic gain can be increased with only limited reduction of the short term commercial genetic gain.

## INTRODUCTION

Successful breeding requires strategies that balance immediate genetic gain with population diversity to sustain long term progress (Jannink 2010). At each selection cycle, plant breeders are facing the choice of new parental lines and the way in which these are mated to improve the mean population performance and generate the genetic variation on which selection will act. Although breeders attempt to account for all available information on candidates, some crosses do not yield selected progeny and do not contribute to genetic gain (Heslot *et al*. 2015). As breeding programs from different companies compete for short term gain, breeders tend to use intensively the most performant individuals sometimes at the expense of genetic diversity (Rauf *et al*. 2010; Gerke *et al*. 2015; Allier *et al*. 2019a). The identification of the crossing plan that maximizes the performance in progeny and limits diversity reduction for long term genetic gain is essential.

Historically, breeders selected the best individuals based on phenotypic observations as a proxy of their breeding value, i.e. the expected value of their progeny. In order to better estimate the breeding value of individuals, phenotypic selection has been complemented by pedigree based prediction of breeding values (Henderson 1984; Piepho *et al*. 2008) and more recently, with cheap high density genotyping becoming available, by genomic prediction of breeding values (Meuwissen *et al*. 2001). In genomic selection (GS), a model calibrated on phenotype and genotype information of a training population is used to predict genomic estimated breeding values (GEBVs) from genome-wide marker information. A truncation selection is commonly applied on GEBVs and the selected individuals are intercrossed to create the next generation. One of the interests of GS is attributed to the acceleration of selection progress by shortening generation interval, increasing selection intensity and accuracy (Hayes *et al*. 2010; Daetwyler *et al*. 2013; Heslot *et al*. 2015). As a consequence, compared to phenotypic selection, GS is expected to accelerate the loss of genetic diversity due to the rapid fixation of large effect regions, but also likely due to the higher probability to select the closest individuals to the training population that are more accurately predicted (Clark et al. 2011; Pszczola et al. 2012). As a result, it has been shown in an experimental study (Rutkoski *et al*. 2015) and by stochastic simulations (Jannink 2010; Lin *et al*. 2016) that GS increases the loss of diversity compared to phenotypic selection. Thus, the optimization of mating strategies in GS breeding programs is a critical area of theoretical and applied research.

Several approaches have been suggested to balance the short and long term genetic gain while selecting crosses using GS. In line with Kinghorn (2011), Pryce et al. (2012), and Akdemir and Sánchez (2016), the selection of a set of crosses, e.g. a list of biparental crosses, requires two components: (i) a cross selection index (CSI) that measures the interest of a set of crosses and (ii) an algorithm to find the set of crosses that maximizes the CSI.

The CSI may consider crosses individually, i.e. the interest of a cross does not depend on the other crosses in the selected set. In classical recurrent GS, candidates with the highest GEBVs are selected and inter-crossed to maximize the expected progeny mean in the next generation. In this case, the CSI is simply the mean of parental GEBVs. However, such an approach neither maximizes the expected response to selection in the progeny, which involves genetic variance generated by Mendelian segregation within each family, nor the long term genetic gain. Alternative measures of the interest of a cross have been suggested to account for parent complementarity, i.e. within cross variability and expected response to selection. Daetwyler et al. (2015) proposed the optimal haploid value (OHV) that accounts for the complementarity between parents of a cross on predefined haplotype segments. Using stochastic simulations, the authors observed that OHV selection yielded higher long term genetic gain and preserved greater amount of genetic diversity than truncated GS. However, OHV does neither account for the position of quantitative trait loci (QTLs) nor the linkage disequilibrium between QTLs (Lehermeier *et al*. 2017b; Müller *et al*. 2018). Schnell and Utz (1975) proposed the usefulness criterion (UC) of a cross to evaluate the expected response to selection in the progeny of the cross. The UC of a cross accounts for the progeny mean (*μ*) that is the mean of parental GEBVs and the progeny standard deviation (*σ*), the selection intensity (*i*) and the selection accuracy (*h*): *UC* = *μ* + *i h σ*. Zhong and Jannink (2007) proposed to predict progeny variance using estimated QTL effects accounting for linkage between loci. Genome-wide marker effects and computationally intensive stochastic simulations of progeny have also been considered to predict the progeny variance (e.g. Mohammadi *et al*. 2015). Recently, an unbiased predictor of progeny variance (*σ*^2^) has been derived in Lehermeier et al. (2017b) for two-way crosses and extended in Allier et al. (2019b) for multi-parental crosses implying up to four parents. Lehermeier et al. (2017b) observed that using UC as a cross selection index increased the short term genetic gain compared to evaluate crosses based on OHV or mean parental GEBV. Similar results have been obtained by simulations in Müller et al. (2018) considering the expected maximum haploid breeding value (EMBV) that is akin to the UC for normally distributed and fully additive traits.

Alternatively, one can consider a more holistic CSI that accounts for the interdependence of crosses in the sense that the interest of a cross depends on the other selected crosses. This is the case in optimal contribution selection, where a set of candidate parents is evaluated as a whole regarding the expected short term gain and the associated risk on loosing long term gain. Optimal contribution selection aims at identifying the optimal contributions (***c***) of candidate parents to the next generation obtained by random mating, in order to maximize the expected genetic value in the progeny (*V*) under a certain constraint on inbreeding (*D*) (Wray and Goddard 1994; Meuwissen 1997; Woolliams *et al*. 2015). Optimal cross selection, further referred as OCS, is an extension of the optimal contribution selection to deliver a crossing plan that maximizes *V* under the constraint *D* by considering additional constraints on the allocation of mates in crosses (Kinghorn *et al*. 2009; Kinghorn 2011; Akdemir and Sánchez 2016; Gorjanc *et al*. 2018; Akdemir *et al*. 2018). In the era of genomic selection, the expected genetic value in progeny (*V*) to be maximized is defined as the mean of parental GEBV (***a***) weighted by parental contributions ***c***, i.e. ***c′a***, and the constraint on inbreeding (*D*) to be minimized is ***c′Kc*** with ***K*** a genomic coancestry matrix. To obtain optimal solutions for the vector of contributions ***c*** and the crossing plan, i.e. pairing of candidates, differential evolutionary algorithms have been suggested (Storn and Price 1997; Kinghorn *et al*. 2009; Kinghorn 2011). Using the concept of optimal contribution selection for mating decisions is common in animal breeding (Woolliams *et al*. 2015) and is increasingly adopted in plant breeding (Akdemir and Sánchez 2016; De Beukelaer *et al*. 2017; Lin *et al*. 2017; Gorjanc *et al*. 2018; Akdemir *et al*. 2018).

In plant breeding one typically has larger biparental families than in animal breeding and especially with GS, the selection intensity within family can be largely increased so that plant breeders much more capitalize on the segregation variance within families compared to animal breeders. In previous works, the genetic gain (*V*) and constraint (*D*) have been defined at the level of the progeny before within family selection. Exceptions are represented by the work of Shepherd and Kinghorn (1998) and Akdemir et al. (2016; 2018) who added a term to *V* accounting for within cross variance assuming LE between QTLs. However, to our knowledge no previous study allowed for linkage disequilibrium (LD) between QTLs. Furthermore, as observed in historical wheat data (Fradgley *et al*. 2019) and using simulations in a maize context (Allier *et al*. 2019b), within family selection also affects the effective contribution of parents to the next generation. This likely biases the prediction of inbreeding/diversity in the next generation, which to our knowledge has not been considered in previous studies.

In this study, we suggest to adjust *V* and *D* terms so that within family selection and the fact that only the best progeny of each family serve as candidates for the next generation are taken into account. We propose to use the usefulness criterion parental contribution (UCPC) approach (Allier *et al*. 2019b) that enables to predict the expected mean performance of the selected fraction of progeny, and to predict the contribution of parents to the selected fraction of progeny. We compared our OCS strategy based on UCPC to account for within family selection with other cross selection strategies, in a long term simulated recurrent genomic selection breeding program involving overlapping generations (Fig. 1A). Our objectives were to demonstrate (1) the interest of UCPC to predict the genetic diversity in the selected fraction of progeny and (2) the interest of accounting for within family selection in OCS for both, short and long term genetic gains.

**Figure 1.**
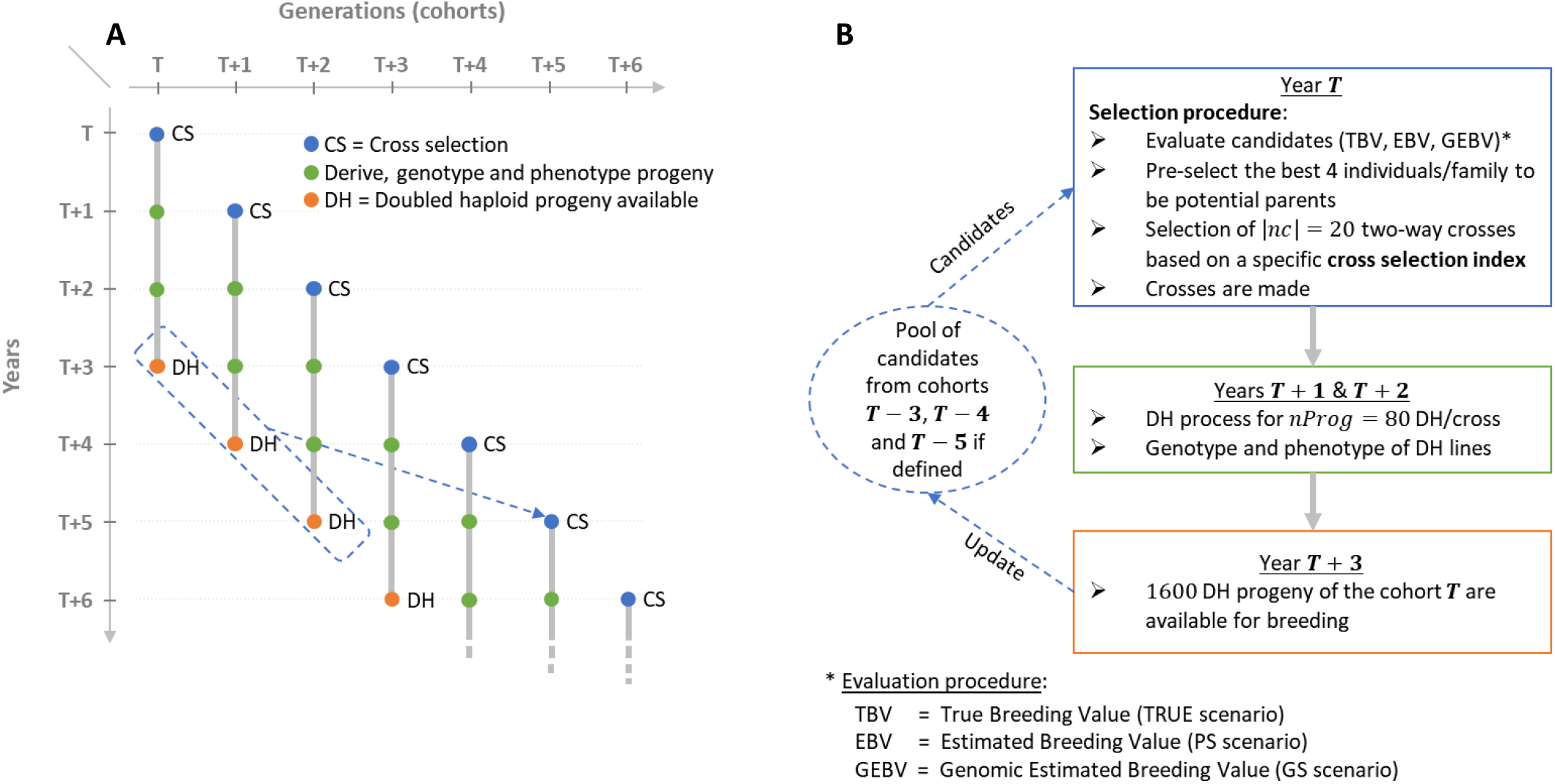
Schematic view of the simulated breeding program: (A) overall view of the breeding program and overlapping cohorts, (B) life cycle of a given post burn-in cohort ***T*** depending on the scenario considered (TRUE with 1,000 known QTL effects, PS in absence of genomic information or GS with 2,000 non causal SNPs estimated effects).

## MATERIAL AND METHODS

### Simulated breeding program

#### Breeding program

We simulated a breeding program to compare the effect of different cross selection indices (CSI) on short and long term genetic gain in a realistic breeding context considering overlapping and connected generations (i.e. cohorts) of three years (Fig. 1A). A detailed description of the simulated breeding program and the material is provided in Supplementary Material (File S1).

Each simulation replicate started from a population of 40 founders sampled among 57 Iodent maize genotypes from the Amaizing project (Rio *et al*. 2019; Allier *et al*. 2019b). We sampled 1,000 biallelic QTLs among 40,478 high-quality single nucleotide polymorphisms (SNPs) from the Illumina MaizeSNP50 BeadChip (Ganal *et al*. 2011) with consensus genetic positions (Giraud *et al*. 2014). The sampling process obeyed two constrains: a QTL minor allele frequency ≥ 0.2 and a distance between two consecutive QTLs ≥ 0.2 cM. Each QTL was assigned an additive effect sampled from a Gaussian distribution with a mean of zero and a variance of 0.05 and the favorable allele was attributed at random to one of the two SNP alleles. We initiated a virtual breeding program starting from the founder genotypes with a burn-in period of 20 years that mimicked recurrent phenotypic selection using doubled haploid (DH) technology. At each generation, phenotypes were simulated considering an error variance corresponding to a trait repeatability of 0.4 in the founder population and no genotype by environment interactions. For phenotyping, every individual was evaluated in four environments in one year. After 20 years of burn-in, we compared different cross selection indices (CSI) for 60 years of recurrent genomic selection using DH technology. Each year, a cohort *T* was generated by 20 two-way crosses (|*nc*| = 20) of 80 DH progeny each (*nProg* = 80). We assumed that three years were needed to produce DH from two-way crosses, and to genotype and phenotype them. Candidate parents of cohort *T* were selected from the available DH of the three cohorts: *T* − 3, *T* − 4 and *T* − 5 (Fig. 1A-B). Per family, the 4 DH lines (i.e. 5%) with the largest breeding values, detailed in “Evaluation scenario” section, were considered as potential parents, yielding 4 DH lines/family x 20 families/cohort x 3 cohorts = 240 potential parents. Considering these *N* = 240 parents, *N*(*N* − 1)/2 = 28,840 two-way crosses are possible. The set of |*nc*| = 20 two-way crosses among these 28,680 candidate crosses was defined using different CSI detailed in the following sections. This simulated scheme yielded overlapping and connected cohorts as it is standard in practical plant breeding (Fig. 1A). Note that 60 years post burn-in corresponded in the simulated context to 20 equivalent non-overlapping generations.

#### Evaluation scenarios

We considered different scenarios for genome-wide marker effects and progeny evaluation. In order to compare several CSI and not blur the comparison between CSI with the uncertainty in marker effect estimates, we mainly focused on the use of the 1,000 known QTL effects and positions (referred to as TRUE scenario). For a representative subset of the CSI, we also considered a more realistic scenario where the effects of 2,000 randomly sampled non causal SNPs were obtained from a G-BLUP model with back solving (Wang *et al*. 2012). This scenario was referred to as GS and marker effects used to predict the CSI were estimated every year with all candidate parents that were phenotyped and genotyped. The progeny were selected on their genomic estimated breeding values (GEBV) considering their phenotypes and genotypes at non causal SNPs. As a benchmark we also considered a phenotypic selection scenario where progeny were selected based on their phenotypic mean (PS). For details on the evaluation models see File S1.

### Cross selection strategies

#### Optimal cross selection not accounting for within family selection

Considering *N* homozygote candidate parents, *N*(*N* − 1)/2 two-way crosses are possible. We define a crossing plan ***nc*** as a set of |*nc*| crosses out of possible two-way crosses, giving the index of selected crosses, i.e. with the *i^th^* element *nc*(*i*) ∈ [1, *N*(*N* − 1)/2]. The (*N* × 1)-dimensional vector of candidate parents contributions ***c*** is defined as:

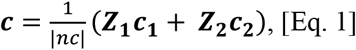

where ***Z*_1_** (respectively ***Z*_2_**) is a (*N* × |*nc*|)- dimensional design matrix that links each *N* candidate parent to the first (respectively second) parent in the set of crosses ***nc, c*_1_** (respectively ***c*_2_**) is a (|*nc*| × 1)-dimensional vector containing the contributions of the first (respectively second) parent to progeny, i.e. a vector of 0.5 when assuming no selection within crosses.

The (*N* × 1)-dimensional vector of candidate parents true breeding values is ***a*** = ***Xβ***_*T*_, where ***X*** = (***x***_1_,…, ***x***_*N*_)′ is the (*N* × *m*)-dimensional matrix of known parental genotypes at *m* biallelic QTLs, where ***x***_*p*_ denotes the (*m* × 1)-dimensional genotype vector of parent *p* ∈ [1, *N*], with the *j^th^* element coded as 1 or −1 for the genotypes AA or aa at QTL *j. **β**_T_* is the (*m* × 1)-dimensional vector of known additive QTL effects for the quantitative agronomic performance trait considered. The genetic gain *V*(***nc***) for this set of two-way crosses is defined as the expected mean performance in the DH progeny:

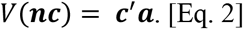

We define the constraint on diversity (*D*) as the mean expected genetic diversity in DH progeny (He, Nei 1973):

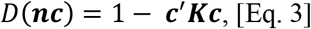

where 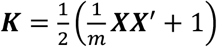 is the (*N* × *N*)-dimensional identity by state (IBS) coancestry matrix between the *N* candidates. File S2 details the relationship between the IBS coancestry among parents (***K***), the parental contributions to progeny (***c***) and the mean expected heterozygosity in progeny 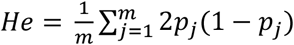 where *p_j_* is the frequency of the genotypes AA at QTL *j* in the progeny.

#### Accounting for within family selection in OCS

In the OCS, as just defined and also considered in Gorjanc et al. (2018), the progeny derived from the ***nc*** crosses are all expected to contribute to the next generation. We suggest to consider *V*(***nc***) and *D*(***nc***) terms accounting for the fact that only a selected fraction of each family will be candidate for the next generation (e.g. 5% per family in our simulation study). For this, we apply the UCPC approach proposed by Allier et al. (2019b) for two-way crosses and extend its use to evaluate the interest of a set ***nc*** of two-way crosses after selection in progeny.

##### UCPC for two-way crosses

Two inbred lines *P*_1_ and *P*_2_ are considered as parental lines for a candidate cross *P*_1_ × *P*_2_ and (***x***_1_, ***x***_2_)′ denotes their genotyping matrix. Following Lehermeier et al. (2017b), the DH progeny mean and progeny variance for the performance trait in the progeny before selection can be computed as:

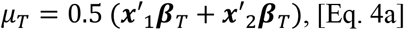

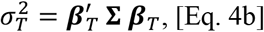

where ***x***_1_, ***x***_2_ and ***β***_*T*_ were defined previously and **Σ** is the (*m* × *m*)-dimensional variance covariance matrix of QTL genotypes in DH progeny defined in Lehermeier et al. (2017b).

To follow parental contributions, we consider *P*_1_ parental contribution as a normally distributed trait (Allier *et al*. 2019b). As we only consider two-way crosses and biallelic QTLs, we can simplify for computational reasons the formulas by using identity by state (IBS) parental contributions computed for polymorphic QTLs between *P*_1_ and *P*_2_ instead of using identity by descent (IBD) parental contributions (Allier *et al*. 2019b). We define the (*m* × 1)-dimensional vector ***β***_*C*1_ to follow *P*_1_ IBS genome contribution at QTLs as 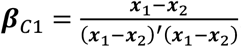. We compute the *P*_1_ mean contribution in the progeny before selection ***μ***_*C*1_ = 0.5 (***x***′_1_***β***_*C*1_ + ***x***′_2_***β***_*C*1_ + 1), where ***x***′_*p*_***β***_*C*1_ + 0.5 is the contribution of *P*_1_ to parent *p*. The progeny variance 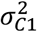 for the *P*_1_ contribution trait in the progeny before selection is computed using Eq. 4b by replacing ***β***_*T*_ by ***β***_*C*1_. The progeny mean for *P*_2_ contribution is then defined as *μ*_*C*2_ = 1 − *μ*_*C*1_.

Following Allier et al. (2019b), we compute the covariance between the performance trait and *P*_1_ contribution trait in progeny as:

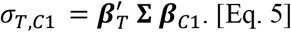

The expected mean performance of the selected fraction of progeny, i.e. usefulness criterion (Schnell and Utz 1975), of the cross *P*_1_ × *P*_2_ is:

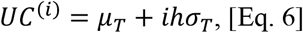

where *i* is the within family selection intensity and the exponent (*i*) in *UC*^(*i*)^ expresses the dependency of *UC* on the selection intensity *i*. We considered a selection accuracy *h* = 1 as in Zhong and Jannink (2007), which holds when selecting on true breeding values. The correlated responses to selection on *P*_1_ and *P*_2_ genome contributions in the selected fraction of progeny are (Falconer and Mackay 1996):

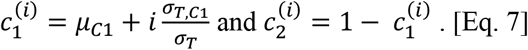

##### Cross selection based on UCPC

Accounting for within family selection intensity *i*, the genetic gain term *V*^(*i*)^(***nc***) for a set of two-way crosses ***nc*** is defined as the expected performance in the selected fraction of progeny:

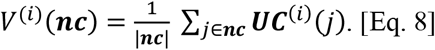

The constraint on diversity *D*^(*i*)^(***nc***) in the selected progeny is defined as:

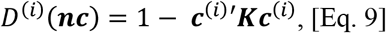

where ***c*^(*i*)^** is defined like ***c*** in Eq. 1 but accounting for within family selection by replacing the ante-selection parental contributions ***c*_1_** and ***c*_2_** by the post-selection parental contributions 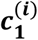 and 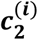 (Eq. 7), respectively. Note that considering the absence of selection in progeny, i.e. *i* = 0, yields *V*^(*i*=0)^(***nc***) being the mean of parent breeding values (Eq. 2) and *D*^(*i*=0)^(***nc***) the expected diversity in progeny before selection (Eq. 3), which is equivalent to optimal cross selection as suggested by Gorjanc *et al*. (2018). The R code (R Core Team 2017) to evaluate a set of crosses as presented in the UCPC based optimal cross selection is provided in File S3.

#### Multi-objective optimization framework

In practice one does not evaluate only one set of crosses but several ones in order to find the optimal set of crosses to reach a specified target that is a function of *V*^(*i*)^(***nc***) and *D*^(*i*)^(***nc***). We use the ***ε***-constraint method (Haimes *et al*. 1971; Gorjanc and Hickey 2018) to solve the multi-objective optimization problem:

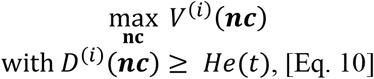

where *He*(*t*), ∀ *t* ∈ [0, *t*\*] is the minimal diversity constraint at time *t*. A differential evolutionary algorithm was implemented to find the set of ***nc*** crosses that is a Pareto-optimal solution of Eq. 10 (Storn and Price 1997; Kinghorn *et al*. 2009; Kinghorn 2011). The direct consideration of *He*(*t*) in the optimization allows to control the decrease in genetic diversity similarly to what was suggested for controlling inbreeding rate in animal breeding (Woolliams *et al*. 1998, 2015). The loss of diversity along time is controlled by the targeted diversity trajectory, i.e. *He*(*t*), ∀ *t* ∈ [0, *t*\*] where *t*\* ∈ ℕ* is the time horizon when the genetic diversity *He*(*t*\*) = *He*\* should be reached. In this study *He*(*t*) is defined as:

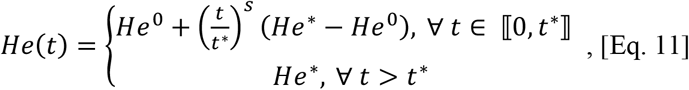

where *He*^0^ is the initial diversity at *t* = 0 and *s* a shape parameter with *s* = 1 for a linear trajectory. Fig. 2 gives an illustration of alternative trajectories that can be defined using Eq. 11.

**Figure 2.**
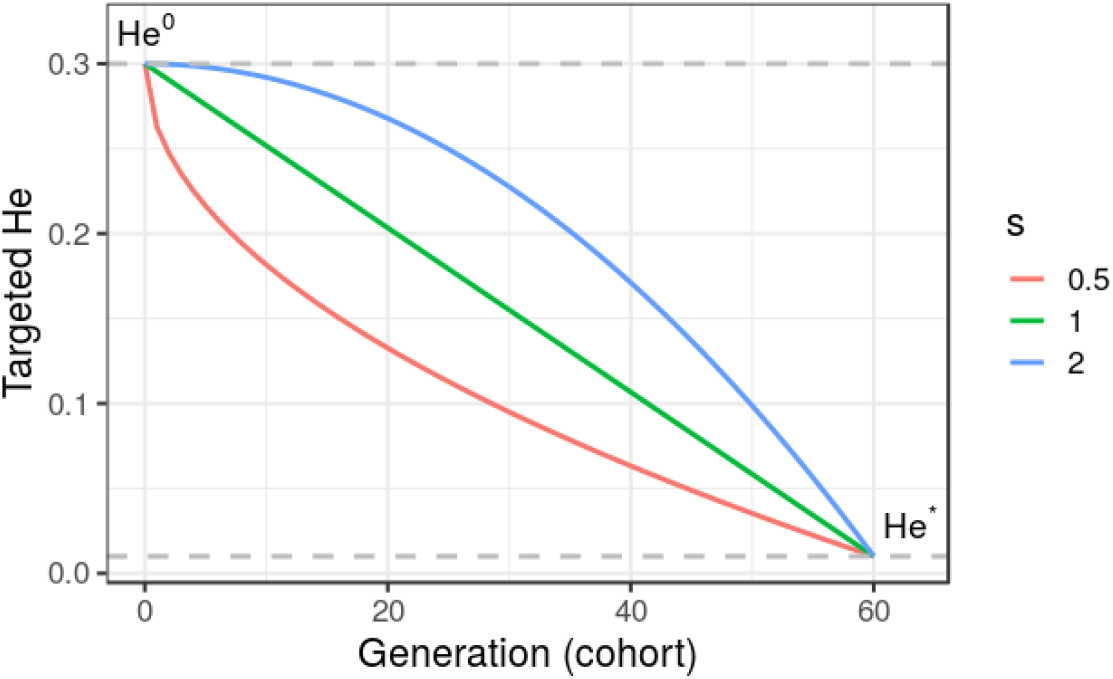
Targeted diversity trajectories for three different shape parameters (s = 1, linear trajectory; s = 2, quadratic trajectory and s = 0.5 inverse quadratic trajectory) for fixed initial diversity (He^0^ = 0.3) at generation 0 and targeted diversity (He* = 0.01) at generation 60 (t* = 60). We considered in this study only linear trajectories (s = 1).

#### Cross selection indices

We considered different cross selection approaches varying in the within family selection intensity (*i*) in *V*^(*i*)^(***nc***), *D*^(*i*)^(***nc***) (Eq. 10) and of the targeted diversity trajectory *He*(*t*) (Eq. 11). We first considered as a benchmark the absence of constraint *D*^(*i*)^(***nc***), i.e. the absence of a diversity trajectory to follow (*He*(*t*) = 0,∀*t*). We defined the cross selection indices PM and UC, respectively considering *V*^(*i*=0)^(***nc***) and *V*^(*i*=206)^(***nc***) with *i* = 2.06 corresponding to select the 5% most performant progeny per family. PM is equivalent to cross the best candidates together without accounting for within cross variance while UC is defined as crossing candidates based on the expected mean performance of the 5% selected fraction of progeny. Notice that the absence of constraint on diversity also means the absence of constraint on parental contributions. To compare optimal cross selection accounting or not for within family selection, we considered three linear diversity trajectories (Eq. 11) with *He*\* = {0.01,0.10,0.15} that should be reached in *t*\* = 60 years. We defined the OCS methods, further referred to as OCS-He*, with *V*^(*i*=0)^(***nc***) and *D*^(*i*=0)^(***nc***). We defined the UCPC cross selection methods, further referred as UCPC-He*, with *V*^(*i*=2.06)^(***nc***) and *D*^(*i*=2.06)^(***nc***). The eight cross selection indices considered are summarized in Table 1.

**Table 1.** Summary of tested cross selection indices (CSI) defined for a set of crosses ***nc*** depending on the within family selection intensity *i*.

| Cross selection index<br>(CSI) | Gain term | Diversity term |
| --- | --- | --- |
| PM | $V^{(i=0)}(\mathbf{nc})$ | - |
| OCS-He* (3 different He*) | $V^{(i=0)}(\mathbf{nc})$ | $D^{(i=0)}(\mathbf{nc})$ |
| UC | $V^{(i=2.06)}(\mathbf{nc})$ | - |
| UCPC-He* (3 different He*) | $V^{(i=2.06)}(\mathbf{nc})$ | $D^{(i=2.06)}(\mathbf{nc})$ |

$He^* = \{0.15; 0.10; 0.01\}$ to be reached linearly ( $s = 1$ ) at the end of simulation ( $t^* = 60$ years). $V^{(i=0)}(\mathbf{nc})$ is the averaged parental mean (PM) of crosses in $\mathbf{nc}$ and $V^{(i=2.06)}(\mathbf{nc})$ is the averaged usefulness criterion (UC) of crosses in $\mathbf{nc}$ considering a within family selection intensity of 2.06. $D^{(i=0)}(\mathbf{nc})$ and $D^{(i=2.06)}(\mathbf{nc})$ are the expected genetic diversity in the progeny before and after within family selection, respectively.

#### Simulation 1: Interest of UCPC to predict the diversity in the selected fraction of progeny

Simulation 1 aimed at evaluating the interest to account for the effect of selection on parental contributions, i.e. post-selection parental contributions (using UCPC), compared to ignore selection, i.e. ante-selection parental contributions (similarly as in OCS), to predict the genetic diversity (He) in the selected fraction of progeny of a set of 20 crosses (using Eq. 9 and Eq. 3, respectively). We considered a within family selection intensity corresponding to selecting the 5% most performant progeny. We used the same genotypes, genetic map and known QTL effects as for the first simulation replicate of the PM cross selection index in the TRUE scenario (Table 1). We extracted the simulated genotypes of 240 DH candidate parents of the first post burn-in cohort (further referred as E1) and of 240 DH candidate parents of the 20th post burn-in cohort (further referred as E2). Due to the selection process, E1 showed a higher diversity and lower performance compared to E2. We randomly generated 300 sets of 20 two-way crosses: 100 sets of intra-generation E1 crosses (E1 x E1), 100 sets of intra-generation E2 crosses (E2 x E2) and 100 sets of inter- and intra-generation crosses randomly sampled (E1 x E2, E1 x E1, E2 x E2). We derived 80 DH progeny per cross and predicted the ante- and post-selection parental contributions to evaluate the post-selection genetic diversity (He) for each set of crosses. We estimated the empirical post-selection diversity for each set of crosses and compared predicted and empirical values considering the mean prediction error as the mean of the difference between predicted He and empirical post-selection He, and the prediction accuracy as the squared correlation between predicted He and empirical post-selection He.

#### Simulation 2: Comparison of different cross selection indices

We ran ten independent simulation replicates of all eight CSI summarized in Table 1 for 60 years post burn-in considering known effects at the 1,000 QTLs (TRUE scenario). We also compared in ten independent simulation replicates the CSI: PM, UC, OCS-He* and UCPC-He* with He*=0.01 considering estimated marker effect at the 2,000 SNPs (GS scenario) and PM based only on phenotypic evaluation (PS scenario). We followed several variables on the 80 DH progeny/family x 20 crosses realized every year. At each cohort *T* ∈ [0,60] with *T* = 0 corresponding to the last burn-in cohort, we computed the additive genetic variance as the variance of the 1600 DH progeny true breeding values (TBV): 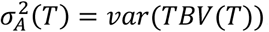). We followed the mean genetic merit of all progeny *μ*(*T*) = *mean*(*TBV*(*T*)) and of the ten most performant progeny 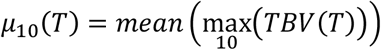 as a proxy of realized performance that could be achieved at a commercial level by releasing these lines as varieties. Then, we centered and scaled the two genetic merits to obtain realized cumulative genetic gains in units of genetic standard deviation at the end of the burn-in (*T* = 0), at the whole progeny level 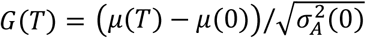 and at the commercial level 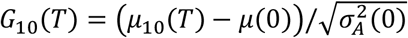.

The interest of long term genetic gain relies on the ability to breed at long term, which depends on the short term economic success of breeding. Following this rationale, we penalized strategies that compromised the short term commercial genetic gain using the weighted cumulative discounted commercial gain following Dekkers et al. (1995) and Chakraborty et al (2002). In practice, we computed the weighted sum of the commercial gain value in each generation 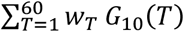, where the weights *w_T_* = 1/(1 + *ρ*)^*T*^, ∀*T* ∈ [1,60] were scaled to have 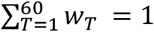 and *ρ* is the interest rate per cohort. For *ρ* = 0, the weights were *w*_*T*∈[1,60]_ = 1/60, i.e. the same importance was given to all cohorts. We compared different values of *ρ* and reported results for *ρ* = 0, *ρ* = 0.04 giving approximatively seven times more weight to short term gain (after 10 years) compared to long term gain (after 60 years) and *ρ* = 0.2 giving nearly no weight to gain reached after 30 years of breeding.

We also measured the genetic diversity as the additive genic variance at QTLs 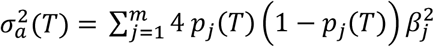, the mean expected heterozygosity at QTLs (He, Nei 1973) 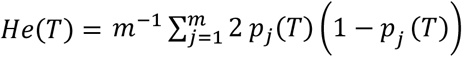, and the number of QTLs where the favorable allele was fixed or lost in the progeny, with *P_j_*(*T*) the allele frequency at QTL *j* ∈ [1, *m*] in the 1600 DH progeny and *β_j_* the additive effect of the QTL *j*. In addition we considered the ratio of additive genetic over genic variance 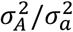 which provides an estimate of the amount of additive genic variance captured by negative covariances between QTL, known as the Bulmer effect under directional selection (Bulmer 1971, 1980; Lynch and Walsh 1999). All these variables were further averaged on the ten simulation replicates.

## RESULTS

### Simulation 1

Compared to the usual approach that ignores the effect of selection on parental contributions, accounting for the effect of within family selection increased the squared correlation (R^2^) between predicted genetic diversity and genetic diversity in the selected fraction of progeny (Fig. 3A-B) for all three types of sets of crosses. The squared correlation between predicted genetic diversity and post-selection genetic diversity for intra-generation sets of crosses was only slightly increased (E1 x E1: from 0.811 to 0.822 and E2 x E2: from 0.880 to 0.888) while the squared correlation for a set of crosses involving also inter-generation crosses was more importantly increased (from 0.937 to 0.987) (Fig. 3A-B). Using post-selection parental contributions instead of ante-selection parental contributions also reduced the mean prediction error (predicted – empirical He) (Fig. 4A-B) for all three types of sets of crosses. The mean prediction error for intra-generation sets of crosses was only slightly reduced (E1 x E1: from 0.006 to 0.005 and E2 x E2: from 0.016 to 0.015) while the mean prediction error for sets involving inter-generation crosses was more reduced (from 0.032 to 0.008) (Fig. 4A-B). The mean prediction error was reduced but still positive when considering post-selection parental contributions, which means that the genetic diversity in the selected fraction of progeny is overestimated. Note that the ante-selection contributions predicted well the empirical genetic diversity before selection for three types of sets of crosses (mean prediction error = 0.000 and R^2^ > 0.992, results not shown).

**Figure 3.**
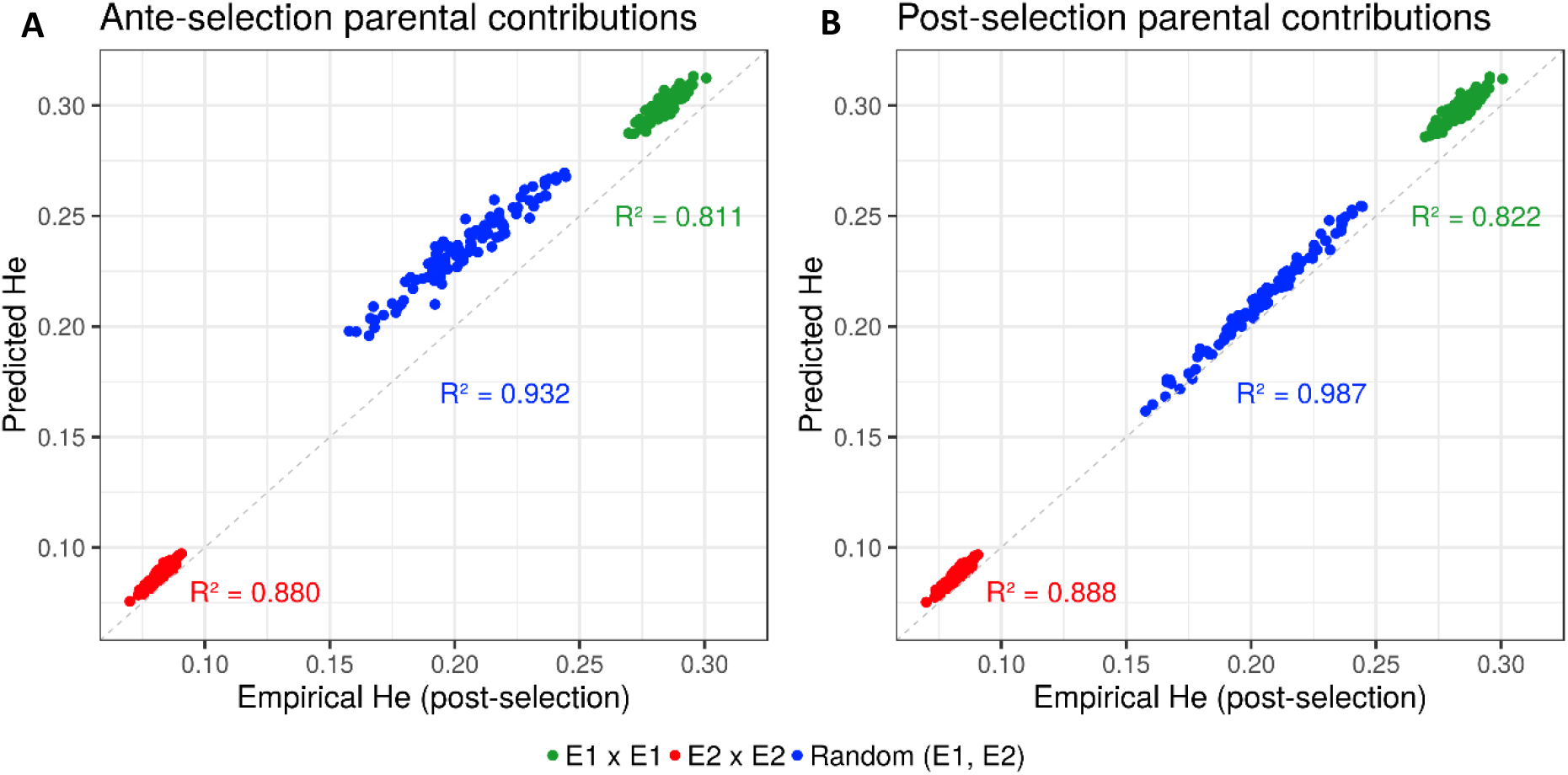
Squared correlations (R^2^) between predicted genetic diversity (He) and empirical He in the selected fraction of progeny of a set of 20 biparental crosses in the TRUE scenario considering (A) ante-selection parental contributions or (B) post-selection parental contributions to predict He. In total 100 sets of each three types of crosses (intra-generation: E1xE1 and E2xE2 or randomly intra and inter-generations: Random (E1,E2)) are shown and the squared correlations between predicted and empirical post-selection He are given in the corresponding color.

**Figure 4.**
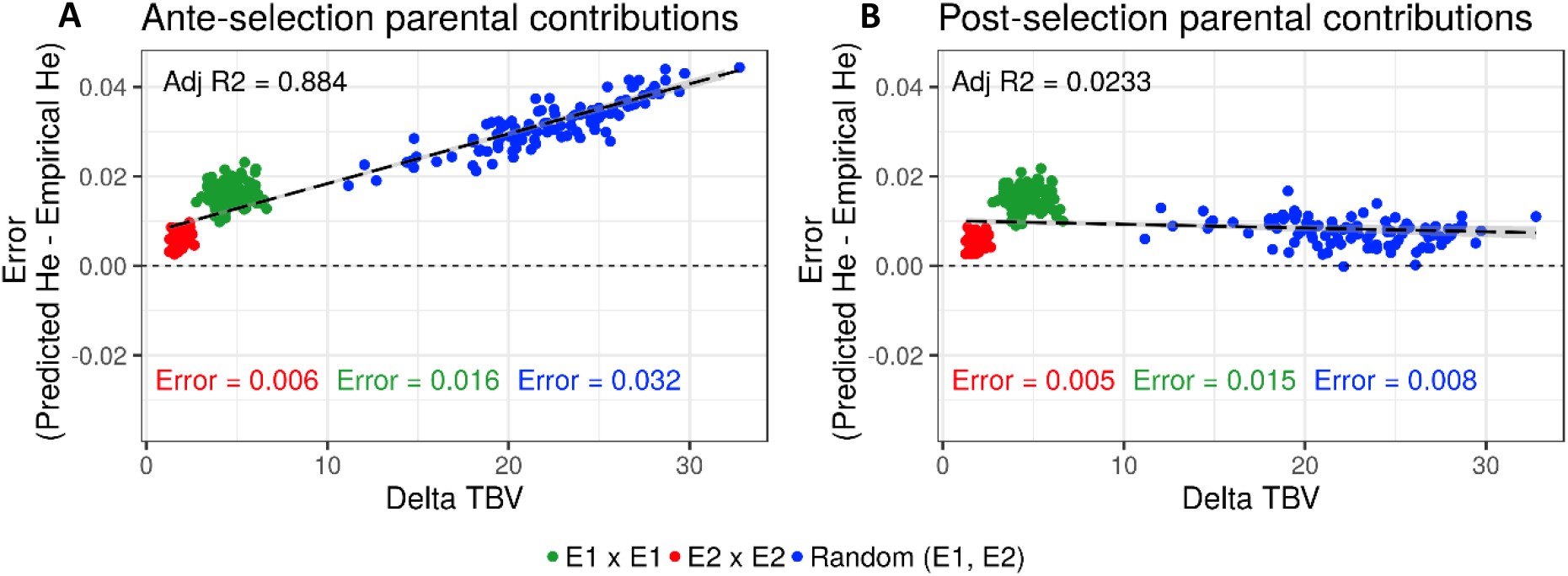
Mean prediction error (predicted - empirical) of predicting the genetic diversity (He) in the selected fraction of progeny of a set of 20 biparental crosses in the TRUE scenario depending on the mean difference of performance between parents (Delta TBV). Mean prediction error is measured as the predicted He - empirical post-selection He, considering (A) ante-selection parental contributions or (B) post-selection parental contributions to predict He. In total 100 sets of each three types of crosses (intra-generation: E1xE1 and E2xE2 or randomly intra and inter-generations: Random (E1,E2)) are shown and the averaged errors are given in the corresponding color.

### Simulation 2

#### Interest of UC over PM

Considering known QTL effects (TRUE scenario), we observed that UC yielded higher short and long term genetic gain at commercial level (G_10_) than PM (9.316 compared to 8.338 ten years post burn-in and 18.293 compared to 15.744 sixty years post burn-in, Fig. 5B-C). When considering the whole progeny mean performance (G), PM outperformed UC for the five first years and after five years UC outperformed PM (Fig. 5A). UC showed higher genic 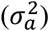 and genetic 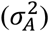 additive variances than PM (Fig. 6A-B) but both yielded a genic and genetic variance near to zero after sixty years of breeding. The genetic over genic variance ratio 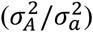 was also higher for UC compared to PM (Fig. 6C). The evolution of genetic diversity (He) along years followed the same tendency as the genic variance (Fig. 7A, Fig. 6A). UC fixed more favorable alleles at QTLs after 60 years (Fig. 7B) and lost less favorable alleles at QTLs than PM in all ten simulation replicates with an average of 243.1 QTLs where the favorable allele was lost compared to 274.9 QTLs for PM (Fig. 7C).

**Figure 5.**
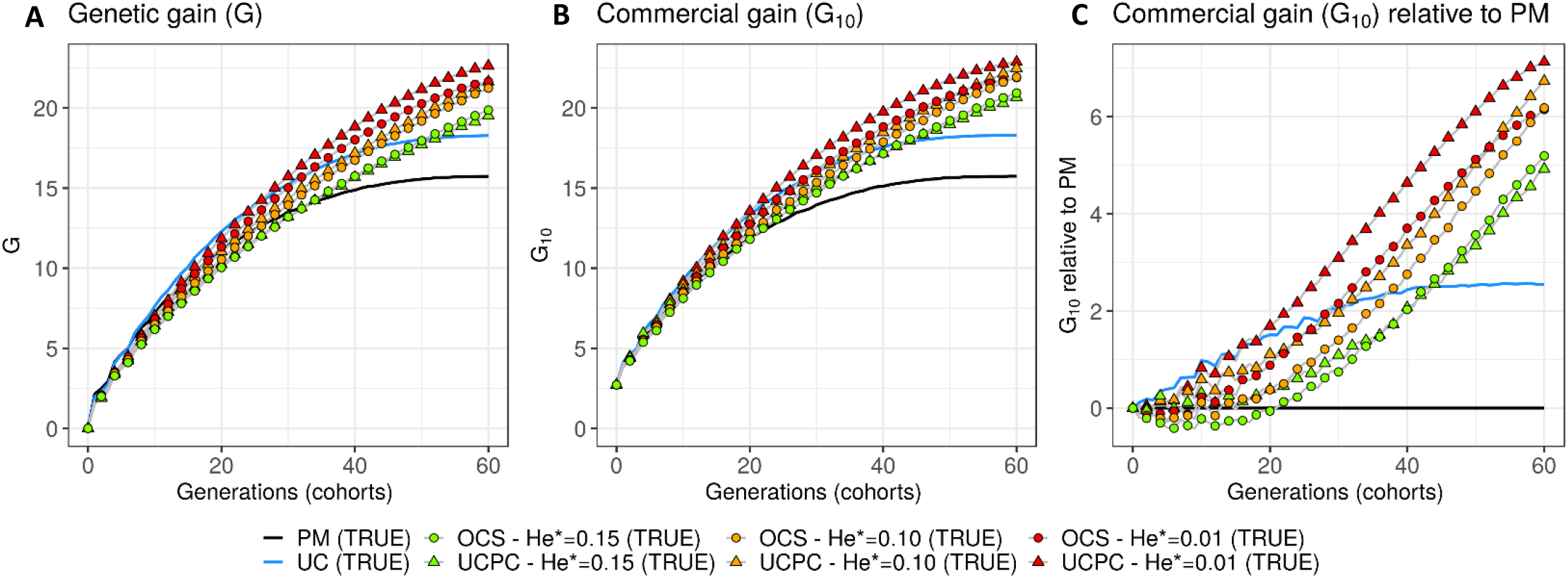
Genetic gains for different cross selection indices in the TRUE scenario (PM: parental mean, UC: usefulness criterion, OCS-He*: optimal cross selection and UCPC-He*: UCPC based optimal cross selection) according to the generations. (A) Genetic gain (G) measured as the mean of the whole progeny, (B) commercial genetic gain (G_10_) measured as the mean of the ten best progeny and (C) G_10_ relative to selection based on parental mean (PM).

**Figure 6.**
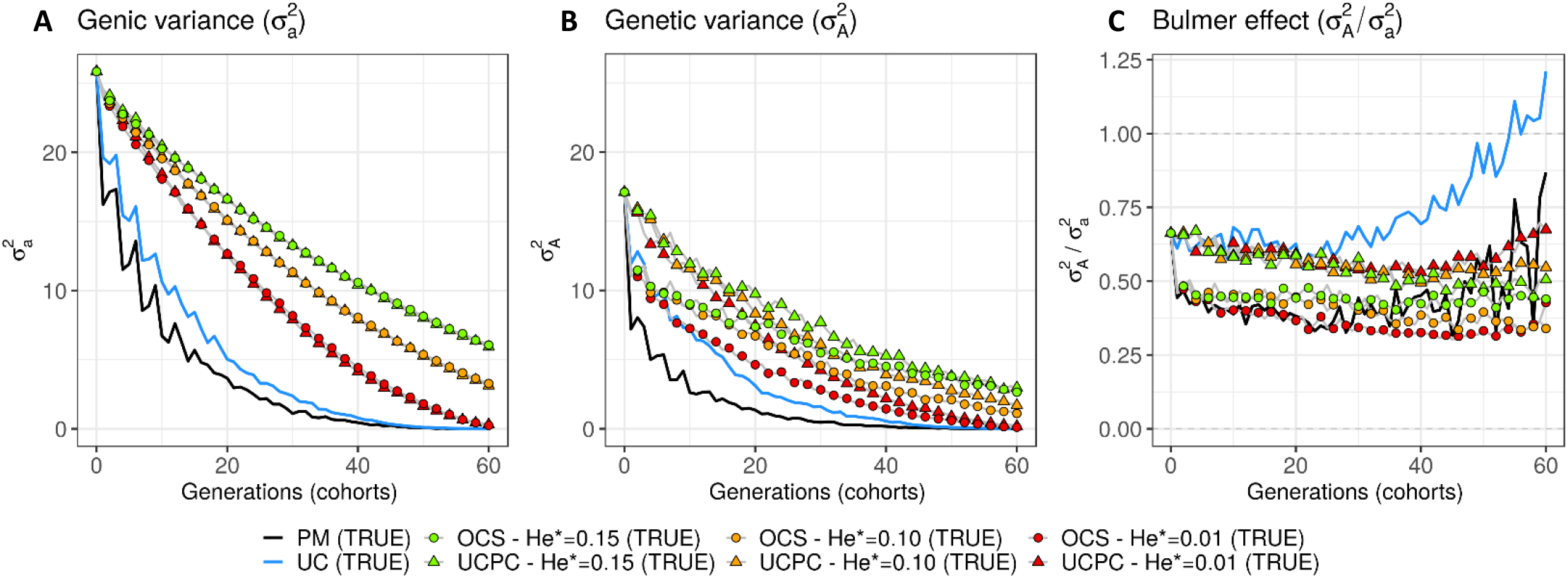
Genetic and genic additive variances for different cross selection indices in the TRUE scenario (PM: parental mean, UC: usefulness criterion, OCS-He*: optimal cross selection and UCPC-He*: UCPC based optimal cross selection) according to the generations. (A) Additive genic variance 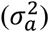 measured on the whole progeny, (B) additive genetic variance 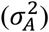 measured on the whole progeny and (C) ratio of genetic over genic variance 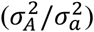 reflecting the Bulmer effect.

**Figure 7.**
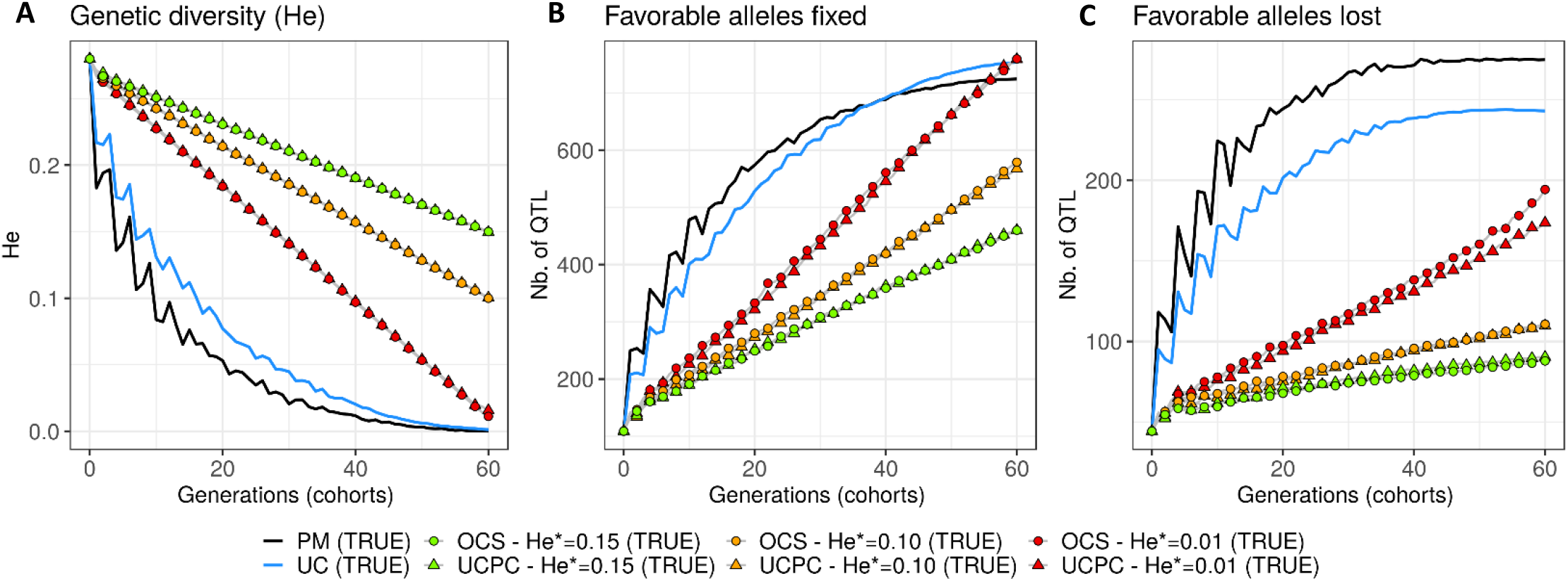
Genetic diversity at QTLs for different cross selection indices in the TRUE scenario (PM: parental mean, UC: usefulness criterion, OCS-He*: optimal cross selection and UCPC-He*: UCPC based optimal cross selection) according to the generations. (A) Genetic diversity at QTLs in the whole progeny (*He*), (B) number of QTLs where the favorable allele is fixed in the whole progeny and (C) number of QTLs where the favorable allele is lost in the whole progeny.

#### Targeted diversity trajectory

Considering known QTL effects (TRUE scenario), the tested optimal cross selection methods OCS-He* and UCPC-He* showed lower short term genetic gain at the whole progeny level (G, Fig. 5A) and at the commercial level (G_10_, Fig. 5B-C) but higher long term genetic gain than UC. The lower the targeted diversity He*, the higher the short and midterm genetic gain at both whole progeny (G, Fig. 5A) and commercial (G_10_, Fig. 5B-C) levels. The higher the targeted diversity He*, the higher the long term genetic gain except for OCS-He*=0.10 and OCS-He*=0.01 that performed similarly after 60 years (on average, G_10_ = 21.925 and 21.892, Fig. 5B). The highest targeted diversity (He* = 0.15) showed a strong penalty at short and midterm, while the intermediate targeted diversity (He* = 0.10) showed a lower penalty at short and midterm compared to the lowest targeted diversity (He* = 0.01) (Fig. 5A-C). For all targeted diversities and all simulation replicates, accounting for within family selection (UCPC-He*) yielded a higher short term commercial genetic gain (G_10_) after 10 years compared to OCS-He* (Fig. 5B-C). Long term commercial genetic gain (G_10_) after 60 years was also higher for UCPC-He* than for OCS-He* with He* = 0.01 in the ten simulation replicates (on average G_10_: 22.869 compared to 21.892) and with He* = 0.10 in nine out of ten replicates (on average G_10_: 22.474 compared to 21.925). However, for He* = 0.15, UCPC-He* outperformed OCS-He* at long term in only three out of ten replicates (on average G_10_: 20.665 compared to 20.938) (Fig. 5B-C). The cumulative commercial gain giving more weight to short term than to long term gain (*ρ* = 0.04) was higher for UCPC-He* than OCS-He* in all simulation replicates for He* = 0.01 (on average, 12.321 compared to 11.675), in all simulation replicates for He*=0.10 (on average, 11.788 compared to 11.278) and in nine out of ten simulation replicates for He*=0.15 (on average, 11.176 compared to 10.884) (Table 2). Cumulative commercial gain giving the same weight to short and long term gain (*ρ* = 0) was also higher for UCPC-He* compared to OCS-He* (Table 2). When giving almost no weight to long term gain after 30 years (*ρ* = 0.2), the best CSI appeared to be UC followed by the UCPC-He* with the lowest constraint on diversity (i.e. low He*).

**Table 2.** Weighted cumulative gain for three different parameters ***ρ*** giving more or less weight to short term gain than to long term gain and assuming known QTL effects (TRUE scenario)

| Cross selection index<br>(CSI) | Weighted cumulative gain |  |  |
| --- | --- | --- | --- |
| | $\rho = 0$<br>(# rank) | $\rho = 0.04$<br>(# rank) | $\rho = 0.2$<br>(# rank) |
| UCPC - He*=0.01 (TRUE) | 15.949 (#1) | 12.321 (#1) | 6.682 (#2) |
| UCPC - He*=0.10 (TRUE) | 15.174 (#2) | 11.788 (#2) | 6.593 (#3) |
| UC (TRUE) | 14.408 (#5) | 11.689 (#3) | 6.822 (#1) |
| OCS - He*=0.01 (TRUE) | 15.148 (#3) | 11.675 (#4) | 6.360 (#5) |
| OCS - He*=0.10 (TRUE) | 14.630 (#4) | 11.278 (#5) | 6.230 (#7) |
| UCPC - He*=0.15 (TRUE) | 14.205 (#6) | 11.176 (#6) | 6.454 (#4) |
| OCS - He*=0.15 (TRUE) | 14.056 (#7) | 10.884 (#7) | 6.103 (#8) |
| PM (TRUE) | 12.609 (#8) | 10.392 (#8) | 6.345 (#6) |

Mean weighted cumulative gain with $\rho = 0$ (constant weight along years), $\rho = 0.04$ (decreasing weight along years) and $\rho = 0.2$ (nearly null weights after 30 years) on the ten independent replicates. For each weighted cumulative gain, the rank of the CSI (# rank) from the most performant (#1) to the less performant (#8) is given.

For a given He* the additive genic variance (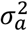, Fig. 6A) and genetic diversity at QTLs (He, Fig. 7A) were constrained by the targeted diversity trajectory for both UCPC-He* or OCS-He*. However, UCPC-He* and OCS-He* behaved differently for genetic variance (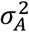, Fig. 6A) resulting in differences for the ratio genetic over genic variances (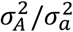, Fig. 6C). UCPC-He* yielded a higher ratio than OCS-He* (Fig. 6C) independently of the targeted diversity He* at short and midterm. For low targeted diversity (He* = 0.01), UCPC-He* showed in all ten replicates a lower number of QTLs where the favorable allele was lost compared to OCS-He* (Fig. 7C, on average 173.6 QTLs-194.3 QTLs).

#### Estimated marker effects

Considering estimated marker effects (GS scenario) yielded lower genetic gain than when considering known marker effects (File S1). However, the short and long term superiority of the usefulness criterion (UC) over the CSI ignoring within cross variance (PM) was consistent with estimated effects (G_10_ = 8.338 compared to 7.713 ten years post burn-in and G_10_ = 15.367 compared to 13.287 sixty years post burn-in, Fig. 8). Similarly, the short and long term superiority of UCPC-He*=0.01 based optimal cross selection over UC and OCS-He*=0.01 was also conserved (G_10_ = 8.162 compared to 7.734 ten years post burn-in and G_10_ = 18.161 compared to 17.528 sixty years post burn-in, Fig. 8). Observations on the genic variance 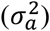 and genetic variance 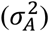 were consistent as well. We also observed that UCPC-He*=0.01 yielded a lower number of QTLs where the favorable allele was lost compared to OCS-He*=0.01 (Fig. 8). PM not considering the marker information, i.e. phenotypic selection (PS scenario), yielded lower short and long term genetic gains than PM considering marker information (G_10_ = 6.402 ten years post burn-in and G_10_ = 10.810 sixty years post burn-in, Fig. 8).

**Figure 8.**
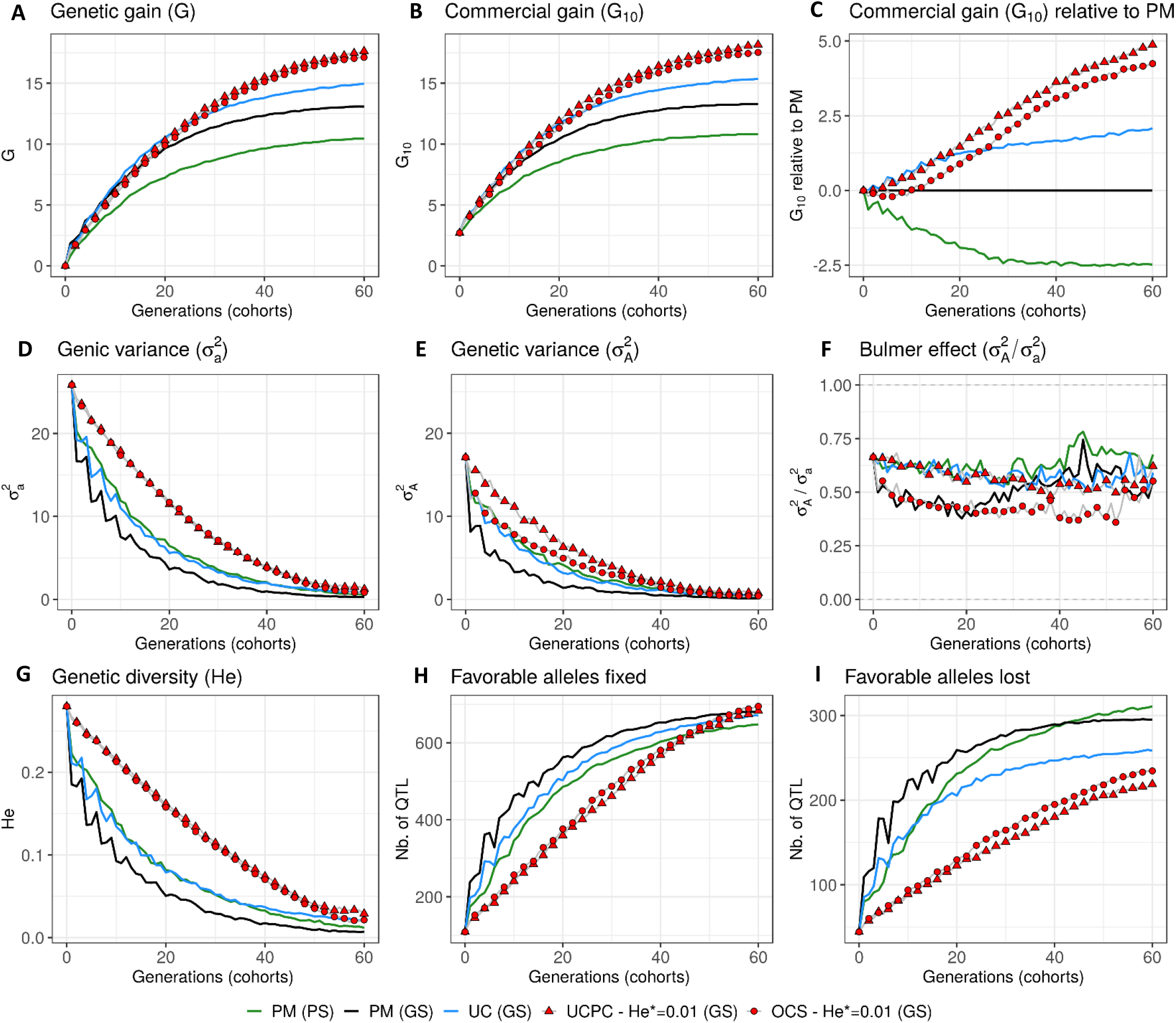
Evolution of different variables for different cross selection indices according to the generations in the GS scenario (PM: parental mean, UC: usefulness criterion, OCS-He*: optimal cross selection and UCPC-He*: UCPC based optimal cross selection for He*=0.01) and in the PS scenario (PM: parental mean). (A) Genetic gain at whole progeny level (G), (B) genetic gain at commercial level (G_10_) and (C) G_10_ relatively to PM (GS), genetic gain is measured on true breeding values. (D) Genic variance at QTLs 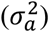, (E) genetic variance of true breeding values 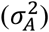 and (F) ratio of genic over genetic variance 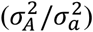, (G) genetic diversity at QTLs and number of QTLs where the favorable allele was fixed (H) and lost (I).

## DISCUSSION

### Predicting the next generation diversity

Accounting for within family selection compared to not accounting for selection to predict the genetic diversity in the selected fraction of progeny increased the squared correlation and reduced the mean error of post-selection genetic diversity prediction (Fig. 4, Fig. 3). The gain in squared correlation (Fig. 3) and the reduction in mean error (Fig. 4), i.e. the interest of UCPC, were more important for parents showing differences in performance. This result is consistent with observations in Allier et al. (2019b) where crosses between two phenotypically distant parents yielded post-selection parental contributions that differ from their expectation before selection (i.e. 0.5). The mean prediction error was always positive, that can be explained by the use in Eq. 9 of genome-wide parental contributions to progeny in lieu of parental contributions at individual QTLs to predict allelic frequency changes due to selection (File S2). As a result, the predicted extreme frequencies at QTLs in the progeny are shrunk towards the mean frequency, leading to an overestimation of the expected heterozygosity (He) (results not shown). Local changes in allele frequency under artificial selection could be predicted following Falconer and Mackay (1996) and Gallais et al. (2007), but this approach would assume linkage equilibrium between QTLs, which is a strong assumption that does not correspond to the highly polygenic trait that we simulated.

### Effect of usefulness criterion on short and long term recurrent selection

In a first approach, we considered no constraint on diversity during cross selection and compared cross selection maximizing the usefulness criterion (UC) or maximizing the parental mean (PM) in the TRUE scenario assuming known QTL effects and positions. The UC yielded higher short term genetic gain at commercial level (G_10_, Fig. 5B-C). This was expected because UC predicts the mean performance of the best fraction of progeny. When considering the genetic gain at the mean progeny level (G, Fig. 5A), UC needed five years to outperform PM. These results underline that UC maximizes the mean performance of the next generation issued from the intercross of selected progeny, sometimes at the expense of the current generation progeny mean performance. This observation is consistent with the fact that candidate parents of the sixth cohort came all from the three first cohorts generated considering UC and thus the sixth cohort took the full advantage of the use of UC (Fig. 1A). This tendency was also observed in simulations by Müller et al. (2018) considering the EMBV approach, akin to the UC for normally distributed additive traits. The UC also showed a higher long term genetic gain at both commercial (G_10_) and whole progeny level (G) compared to intercross the best candidate parents (PM). This long term gain was driven by a higher additive genic variance at QTLs (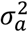, Fig. 6A) and a lower genomic covariance between QTLs (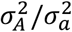, Fig. 6C) resulting in a higher additive genetic variance in UC compared to PM (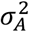, Fig. 6B). Note that with lower 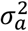 the ratio 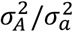 becomes less interpretable at long term (Fig. 6C). UC also better managed the fixation (Fig. 7B) or the maintenance (Fig. 7C) of the favorable allele at QTLs compared to PM. These results highlight the interest of considering within cross variance in cross selection for improving long term genetic gain as observed in Müller et al. (2018).

### Accounting for within family variance in optimal cross selection

Assuming known marker effects, we observed that to consider a constraint on diversity, i.e. in optimal cross selection, always maximized the long term genetic gain along with a variable penalty at short term gain compared to no constraint on diversity when selecting crosses (e.g. UC). We further compared the OCS (Gorjanc *et al*. 2018) with the UCPC based optimal cross selection that accounts for the fact that only a selected fraction of each family is candidate for the next generation. In the optimization framework considered, we compared the ability of UCPC (referred to as UCPC-He*) and OCS (referred to as OCS-He*) to convert a determined loss of diversity into genetic gain. For a given diversity trajectory, UCPC-He* yielded higher short term commercial gain than OCS-He*. Both, OCS-He* and UCPC-He* yielded similar additive genic variance 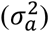 but we observed differences in terms of the ratio 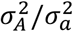. As expected under directional selection, the ratio 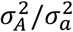 was positive and inferior to one, revealing a negative genomic covariance between QTLs (Bulmer 1971). UCPC-He* yielded a higher ratio, i.e. lower repulsion, and thus a higher additive genetic variance 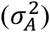 than OCS-He* for a similar He*. This explains the higher long term genetic gain at commercial and whole progeny levels observed for UCPC-He*. This result supports the idea, suggested in Allier et al. (2019a), that accounting for complementarity between parents when defining crossing plans is an efficient way to favor recombination events to reveal part of the additive genic variance hidden by repulsion between QTLs. For low targeted diversity (He* = 0.01), UCPC-He* also appeared to better manage the rare favorable alleles at QTLs than OCS-He*. These results highlighted the interest of UCPC based optimal cross selection to convert the loss of genetic diversity into genetic gain by maintaining more rare favorable alleles and limiting repulsion between QTLs. Note that the superiority of UCPC-He* over OCS-He* for long term genetic gain decreased when considering higher targeted diversity. In case of higher targeted diversity (He* = 0.15), the loss of diversity was likely not sufficient to fully express the additional interest of UCPC compared to OCS to convert diversity into genetic gain. In this case UCPC-He* and OCS-He* performed similarly. Accounting for within cross variance to measure the expected gain of a cross in optimal cross selection was already suggested in Shepherd and Kinghorn (1998). More recently, Akdemir and Sánchez (2016) and Akdemir et al. (2018) accounted for within cross variance considering linkage equilibrium between QTLs. Akdemir and Sánchez (2016) also observed that accounting for within cross variance during cross selection yielded higher long term mean performance with a penalty at short term mean progeny performance.

Short term economic returns condition the ability of a breeder to target long term genetic gain. Hence, it is necessary to make sure that tested breeding strategy do not compromise too much the short term commercial genetic gain. For this reason, we considered the weighted cumulative discounted commercial gain following Dekkers et al. (1995) and Chakraborty et al (2002) as a summary variable to evaluate CSI while giving more or less weight to short and long term performance. UCPC-He* outperformed OCS-He* for a given He* considering either uniform weights (*ρ* = 0) or giving approximately seven time more weight to short term gain compared to long term gain (*ρ* = 0.04). This was also true when focusing only on short term gain (*ρ* = 0.2), but in this case the best model was UC without accounting for diversity while selecting crosses (Table 2).

### Practical implementations in breeding

#### UCPC with estimated marker effects

In simulations, we firstly considered 1,000 QTLs with known additive effects sampled from a centered normal distribution. For a representative subset of cross selection indices (PM, UC, UCPC-He* and OCS-He* with He*=0.01, Fig. 8) we considered 2,000 SNPs estimated effects. The main conclusions were consistent considering both estimated and known marker effects, supporting the practical interest of UCPC based optimal cross selection (Fig. 8). With estimated marker effects instead of known QTL effects, the predicted progeny variance (*σ*^2^) corresponded to the variance of the predicted breeding values which are shrunk compared to true breeding values depending on the model accuracy (referred to as variance of posterior mean, VPM in Lehermeier *et al*. (2017a; b)). An alternative would be to consider the marker effects estimated at each sample of a Monte Carlo Markov Chain process, e.g. using a Bayesian Ridge Regression, to obtain an improved estimate of the additive genetic variance (referred to as posterior mean variance, PMV in Lehermeier *et al*. (2017a; b)).

In practice, QTL effects are unknown, so the selection of progeny cannot be based on true breeding values and thus the selection accuracy (*h*) is smaller than one. In our simulation study assuming unknown QTLs (GS scenario), progeny were selected based on estimated breeding values taking into account genotypic information as well as replicated phenotypic information leading to a high selection accuracy, as it can be encountered in breeding. In order to shorten the cycle length of the breeding scheme, selection of progeny can be based on predicted GEBVs of genotyped but not phenotyped progeny. In such a case, the selection accuracy (*h*) will be considerably reduced. We assume that selection based on UCPC can be improved when using PMV instead of VPM and by taking into account the proper selection accuracy (*h*) within crosses adapted to the selection scheme. When selection is based on predicted values, i.e. genotyped but not phenotyped progeny, the shrunk predictor VPM might present a good approximation of (*hσ*)^2^.

#### UCPC based optimal cross selection

In this study, we assumed fully homozygous parents and two-way crosses. However, neither the optimal cross selection nor UCPC based optimal cross selection are restricted to homozygote parents. Considering heterozygote parents in optimal cross selection is straightforward. Following the extension of UCPC to four-way crosses (Allier *et al*. 2019b), UCPC optimal cross selection can be used for phased heterozygous individuals, as it is commonly the case in perennial plants or animal breeding. We considered an inbred line breeding program but the extension to hybrid breeding is of interest for species as maize. The use of testcross effects, i.e. estimated on hybrids obtained by crossing candidate lines with lines from the opposite heterotic pool, in UCPC based optimal cross selection is straightforward and so the UCPC based optimal cross selection can be used to improve each heterotic pool individually. In order to jointly improve two pools, further investigations are required to include dominance effects in UCPC based optimal cross selection. In addition, this would imply that crossing plans in both pools are jointly optimized to manage genetic diversity within pools and complementarity between pools.

We considered a within family selection intensity corresponding to the selection of the five percent most performant progeny as candidates for the next generation. Equal selection intensities were assumed for all families but in practice due to experimental constraints or optimized resource allocation (e.g. generate more progeny for crosses showing high progeny variance but low progeny mean), within family selection intensity can be variable. Different within family selection intensities (see Eq. 8 and Eq. 9) can be considered in UCPC based optimal cross selection, but an optimization regarding resource allocation of the number of crosses and the selection intensities within crosses warrants further investigations. However, in marker-assisted selection schemes based on QTL detection results (Bernardo *et al*. 2006) an optimization of selection intensities per family was observed to be only of moderate interest.

Proposed UCPC based optimal cross selection was compared to OCS in a targeted diversity trajectory context. We considered a linear trajectory but any genetic diversity trajectory can be considered (e.g. Fig. 2). The optimal diversity trajectory cannot be easily determined and depends on breeding objectives and data considered. Optimal contribution selection in animal breeding considers a similar *ϵ*-constraint optimization with a targeted inbreeding trajectory determined by a fixed annual rate of inbreeding (e.g. 1% advocated by the FAO, Woolliams *et al*. 1998). Woolliams (2015) argued that the optimal inbreeding rate is also not straightforward to define. An alternative formulation of the optimization problem to avoid the use of a fixed constraint is to consider a weighted index (1 − *α*)*V*(***nc***) + *αD*(***nc***), where *α* is the weight balancing the expected gain *V*(***nc***) and constraint *D*(***nc***) (De Beukelaer *et al*. 2017). However, the appropriate choice of *α* is difficult and is not explicit either in terms of expected diversity nor expected gain.

#### Introgression of diversity and anticipation of a changing breeding context

We considered candidate parents coming from the three last overlapping cohorts (Fig. 1) in order to reduce the number of candidate crosses during the progeny covariances prediction (UCPC) and the optimization process. This yielded elite candidate parents that were not directly related (no parent-progeny) and that did not show strong differences in performances, which is standard in a commercial plant breeding program focusing on yield improvement. However, when the genetic diversity in a program is too low so that long term genetic gain is compromised, external genetic resources need to be introgressed by crosses with internal elite parents. As suggested by results of simulation 1, we conjecture that the advantage of UCPC based optimal cross selection over OCS increases in such a context where heterogeneous, i.e. phenotypically distant, genetic material are crossed. This requires investigations that we hope to address in subsequent research.

Our simulations also assumed fixed environments and a single targeted trait over sixty years. However, in a climate change context and with rapidly evolving societal demands for sustainable agricultural practices, environments and breeders objectives will likely change over time. In a multi-trait context, the multi-objective optimization framework proposed in Akdemir et al. (2018) can be adapted to UCPC based optimal cross selection. The upcoming but yet unknown breeding objectives make the necessity to manage genetic diversity even more important than highlighted in this study.

## Author contributions

ST, CL, AC and LM supervised the study. AA performed the simulations and wrote the manuscript. ST worked on the implementation in the simulator. All authors reviewed and approved the manuscript.

## Acknowledgments

This research was funded by RAGT2n and the ANRT CIFRE Grant n° 2016/1281 for AA.

## Supplementary material

File S1 details the simulated breeding program; File S2 demonstrates the relationship between IBS coancestry and genetic diversity (He) in progeny and File S3 provides the R code to evaluate a set of crosses as presented in the UCPC based optimal cross selection.

