## Supplementary material for "Improving short and long term genetic gain by accounting for within family variance in optimal cross selection": File S1

**Additional material**

***Material***

We initiated simulations with the genome of 57 maize Iodent inbred lines (*Zea mays L.*) (Allier *et al.* 2019). These lines were genotyped with the Illumina MaizeSNP50 BeadChip (Ganal *et al.* 2011). After quality control and imputation, 40,478 high-quality SNPs were retained. The genetic map was obtained by predicting genetic positions from physical positions on the reference genome B73-v4 (Jiao *et al.* 2017) using a spline-smoothing interpolating procedure described in Bauer *et al.* (2013) and the dent genetic map in Giraud *et al.* (2014). At each simulation replicate we randomly sampled 40 lines to be the founder population. We randomly sampled 1,000 SNPs to be additive biallelic quantitative trait loci (QTL) of a polygenic trait. The sampling of QTL obeyed two constraints: QTL minor allele frequency $\geq 0.2$ and distance between two consecutive QTL $\geq 0.2$ cM. Each QTL was assigned an additive effect from a Gaussian distribution with a mean of zero and a variance of 0.05. For the scenario where the 1,000 QTLs were unknown, we randomly sampled 2,000 non causal SNPs as genomewide markers used for evaluation (see “Evaluation model” section).

***Simulation scheme***

We aimed at comparing the effect of parent selection and allocation methods on short and long term genetic gains in a realistic breeding context using doubled haploid (DH) technology and considering overlapping and connected cohorts (i.e. generations) of three years as illustrated in Figure 1A. We considered that the process to derive DH lines from a cross and to phenotype and genotype DH lines took three years. Furthermore we considered as candidate parents of a new cohort only the DH progeny of the three last cohorts. For sake of clarity, the candidate parents of cohort $T$ were selected from the available DH progeny of the three cohorts: $T-3$, $T-4$ and $T-5$ (Fig. 1A-B). Within this breeding context, we defined a burn-in period of 20 years starting from founders that mimicked a phenotyping selection (PS) program using DH technology (more details in the “phenotyping” and “evaluation model” sections). Afterward, we compared different cross selection strategies during 60 years of breeding. We considered either that we had access to the 1,000 QTL effects (TRUE scenario) or that we estimated the effects of the 2,000 non causal SNPs (GS scenario). We also considered the absence of genomic information for selection, i.e. phenotypic selection (PS scenario).

We can distinguish the following simulation phases for the cohorts $\boldsymbol{T} \in [\mathbf{1}, \mathbf{80}]$:

- **Burn-in Phase 1 ($\boldsymbol{T} \in [\![\mathbf{1}; \mathbf{3}]\!]$): Initialization**

Every year during the three first years, a cohort was initiated by randomly generating 20 biparental crosses from the 40 founders. We derived 80 DH lines per cross. Note that lines can contribute as parents to different cohorts, so that different cohorts can share the same crosses at this stage.

- **Burn-in Phase 2 ($\boldsymbol{T} \in [\![\mathbf{4}; \mathbf{20}]\!]$)**

The second phase of burn-in mimicked 17 years of phenotypic selection to build up extensive linkage disequilibrium to compare scenarios in a realistic ongoing breeding context. In burn-in phase 2, phenotypic selection (PS) was used to estimate breeding value of candidate lines from the three last cohorts ($T-3$, $T-4$ and $T-5$, if available). After selecting the 4 best DH progeny per family (i.e. 5%), the overall 50 best progeny out of 3 cohorts x 20 families/cohort x 4 DH/family = 240 DH progeny were considered as potential parents of the cohort and were randomly mated to generate 20 biparental families of 80 DH lines. Note that lines can contribute as parents to different cohorts, so that different cohorts can share the same crosses at this stage. Burn-in ended up with overlapping cohorts connected by the pedigree as it can be found in real breeding program.

- **Post burn-in ($\boldsymbol{T} \in [\![\mathbf{21}; \mathbf{70}]\!]$)**

In post burn-in, the life cycle of a cohort was similar to burn-in phase 2 except changes in the way to evaluate, select and allocate parents (Fig. 1B).

***Phenotyping***

For phenotyping, we considered environmental effects sampled in a normal distribution of mean zero and variance 25 and did not consider genotype by environment interactions. Each cohort was evaluated in $N_{loc} = 4$ locations in one year, i.e. four environments. At each simulation replicate, five founder lines were randomly sampled to be check individuals phenotyped every year. Environmental errors were sampled from a normal distribution with mean zero and an error variance $\sigma_\epsilon^2$ defined by the initial repeatability in the founder population $r = \frac{\sigma_G^2}{\sigma_G^2 + \sigma_\epsilon^2} = 0.4$. This led to a heritability in the founder population of $h^2 = \frac{\sigma_G^2}{\sigma_G^2 + \sigma_\epsilon^2 / N_{loc}} = 0.73$. Note that the repeatability and heritability varied along selection cycles relatively to the evolution of additive genetic variance $\sigma_G^2$.

***Evaluation model***

Different evaluation models were considered and should be distinguished at this stage. For phenotypic selection (PS scenario), the phenotypes of progeny were used to estimate their breeding values (EBV). We distinguished two scenarios using genomic information. On one hand, the 1,000 QTL positions and effects were known (TRUE scenario) and the evaluation consisted in summing the individual additive QTL effects to obtain the true breeding value (TBV) of progeny. On the other end, the 1,000 QTL positions and effects were unknown (GS scenario) and 2,000 SNP effects were estimated using the phenotypes and genotypes of the progeny from the three last cohorts. The progeny were selected on their genomic estimated breeding values (GEBV).

The breeding value of progeny (EBV in PS or GEBV in GS) were estimated in Model 1 S1 fitted using mixed model software blup-f 90 (Misztal 2008) with AI-REML variance component estimates:

$$\boldsymbol{Y} = \boldsymbol{1}\mu + \boldsymbol{E}\boldsymbol{\beta}_{\boldsymbol{Env}} + \boldsymbol{W}\boldsymbol{u} + \boldsymbol{\epsilon}, \text{ (Model 1 S1)}$$

where $\boldsymbol{Y}$ is the vector of phenotypic values, $\mu$ is the intercept, $\boldsymbol{E}$ is the incidence matrix for environmental effects, $\boldsymbol{\beta}_{\boldsymbol{Env}}$ is the vector of environmental fixed effects, $\boldsymbol{W}$ is the incidence matrix of individual breeding value random effects $\boldsymbol{u}$, $\boldsymbol{u} \sim N(\boldsymbol{0}, \sigma_G^2\boldsymbol{U})$ is the vector of breeding value random effects with $\sigma_G^2\boldsymbol{U}$ its variance-covariance matrix and $\boldsymbol{\epsilon}$ is the vector of residual random terms $\boldsymbol{\epsilon} \sim N(\boldsymbol{0}, \sigma_\epsilon^2\boldsymbol{I})$ independent and identically distributed. For phenotypic selection (PS), the individuals were assumed independent, i.e. $\boldsymbol{u} \sim N(\boldsymbol{0}, \sigma_G^2 \boldsymbol{I})$. For genomic selection (GS), the covariance between individuals was modeled using the genomic relationship matrix $\boldsymbol{G}$, i.e. $\boldsymbol{u} \sim N(\boldsymbol{0}, \sigma_G^2 \boldsymbol{G})$. Hereby, $\boldsymbol{G}$ was estimated according to VanRaden (2008) using the 2,000 non causal loci:

$$\boldsymbol{G} = \frac{\boldsymbol{Z}\boldsymbol{Z}'}{4\sum_j p_j(1-p_j)}$$

where, $\boldsymbol{Z}$ contains the centered allele counts, with elements computed as $x_{ij} + 1 - 2p_j$, where the element $x_{ij} \in \{-1,1\}$ is the genotype for individual $i$ at non causal locus $j$ and $p_j$ is the frequency of the allele for which the homozygous genotype is coded 1 at non causal locus $j$. Estimated marker effects $\widehat{\boldsymbol{\beta}_{\boldsymbol{T}}}$ were obtained by back-solving: $\widehat{\boldsymbol{\beta}_{\boldsymbol{T}}} = \boldsymbol{Z}'(\boldsymbol{Z}\boldsymbol{Z}')^{-\boldsymbol{1}}\widehat{\boldsymbol{u}}$ (Wang *et al.* 2012) and used in lieu of known QTL effects $\boldsymbol{\beta}_{\boldsymbol{T}}$.

***Simulation of progeny genotypes***

Doubled haploid progeny genotypes were simulated considering meiosis events without crossover interference. The number of chiasmata was drawn from a Poisson distribution with $\lambda$ equal to the chromosome length in Morgan, and crossover positions were determined using the recombination frequency obtained using the Haldane mapping function (Haldane 1919).
