## Supplementary material for "Improving short and long term genetic gain by accounting for within family variance in optimal cross selection": File S2

**Relationship between IBS coancestry and genetic diversity in progeny**

The identity by state (IBS) coancestry between $N$ inbred parents is defined as:

$$\boldsymbol{K} = 0.5\left(\frac{1}{m}\,(\boldsymbol{X}\boldsymbol{X}') + \boldsymbol{1}_N\boldsymbol{1}_N'\right), \text{(Eq. 1)}$$

where, $\boldsymbol{X}$ is the genotyping matrix of the $N$ parents in line and $m$ loci in column, with elements coded -1 or 1 and $\boldsymbol{1}_N$ is a $N$-dimensional column vector of ones.

Considering the $N$-dimensional column vector of expected parental genomewide contributions $\boldsymbol{c}$, with $c(j),\ j \in [1, N]$ the contribution of the parent $j$ to progeny, the mean expected IBS coancestry in progeny is:

$$IBS = \boldsymbol{c}'\boldsymbol{K}\,\boldsymbol{c} = 0.5\left[\frac{1}{m}(\boldsymbol{c}'\boldsymbol{X}\boldsymbol{X}'\boldsymbol{c}) + \boldsymbol{c}'\boldsymbol{1}_N\boldsymbol{1}_N'\boldsymbol{c}\right]. \text{(Eq. 2a)}$$

Note that $\boldsymbol{c}'\boldsymbol{1}_N\boldsymbol{1}_N'\boldsymbol{c} = 1$ since $\sum_{j=1}^{N}\boldsymbol{c}(j) = 1$. Then, Eq. 2a simplifies:

$$IBS = 0.5\left[\frac{1}{m}(\boldsymbol{c}'\boldsymbol{X}\boldsymbol{X}'\boldsymbol{c}) + 1\right] \text{(Eq. 2b)}$$

The mean expected genetic diversity (He) in progeny is:

$$He = \frac{1}{m}\,\boldsymbol{1}_m'\left(2\,\boldsymbol{p}\circ(\boldsymbol{1}_m - \boldsymbol{p})\right), \text{(Eq. 3a)}$$

where $\boldsymbol{1}_m$ is a $m$-dimensional column vector of ones and $\boldsymbol{p}$ is the $m$-dimensional column vector of expected allelic frequencies in progeny:

$$\boldsymbol{p} = 0.5\,((\boldsymbol{X} + \boldsymbol{1}_N\boldsymbol{1}_m')' \circ \mathbf{C})\,\boldsymbol{1}_N, \text{(Eq. 4a)}$$

where $\mathbf{C}$ is the $(m \text{ x } N)$-dimensional matrix of expected local parental contributions to progeny with $C(i,j),\ i \in [1,m],\ j \in [1,N]$ the contribution of parent $j$ to progeny at the locus $i$. $C(i,j), \forall\, i \in [1,m]$

is further approximated by the genomewide parental contribution to progeny $c(j)$. Consequently, the $m$-dimensional column vector of expected allelic frequencies (Eq. 4a) is approximated as:

$$\widetilde{\boldsymbol{p}} = 0.5\ (\boldsymbol{X} + \mathbf{1}_N \mathbf{1}'_m\ )'\boldsymbol{c}. \text{ (Eq. 4b)}$$

We replace $\boldsymbol{p}$ by its approximation $\widetilde{\boldsymbol{p}}$ in Eq. 3a:

$$\widetilde{He} = \frac{1}{m}\ \mathbf{1}'_m\ \left((\boldsymbol{X}'\boldsymbol{c} + \mathbf{1}_m \mathbf{1}'_N \boldsymbol{c}) \circ (\mathbf{1}_m - 0.5\ \boldsymbol{X}'\boldsymbol{c} - 0.5\ \mathbf{1}_m \mathbf{1}'_N \boldsymbol{c})\right). \text{ (Eq. 5a)}$$

Note that $\mathbf{1}_m \mathbf{1}'_N \boldsymbol{c} = \mathbf{1}_m$ and Eq. 5a becomes:

$$\widetilde{He} = \frac{1}{m}\ \mathbf{1}'_m\ \left((\boldsymbol{X}'\boldsymbol{c} + \mathbf{1}_m) \circ (0.5\ \mathbf{1}_m - 0.5\ \boldsymbol{X}'\boldsymbol{c})\right)$$

$$= \frac{1}{m}\ \mathbf{1}'_m\ \left(0.5\ (\mathbf{1}_m - \boldsymbol{X}'\boldsymbol{c} \circ \boldsymbol{X}'\boldsymbol{c})\right)$$

$$= 0.5\ \left(1 - \frac{1}{m}\ \mathbf{1}'_m\ (\boldsymbol{X}'\boldsymbol{c} \circ \boldsymbol{X}'\boldsymbol{c})\right). \text{ (Eq. 5b)}$$

Let us note $\boldsymbol{v} = \boldsymbol{X}'\boldsymbol{c}$. It can be shown that $\mathbf{1}'_m\ (\boldsymbol{v} \circ \boldsymbol{v}) = \ \boldsymbol{v}'\boldsymbol{v}$, resulting in:

$$\widetilde{He} = 0.5\ \left(1 - \frac{1}{m}\ (\boldsymbol{c}'\boldsymbol{X}\boldsymbol{X}'\boldsymbol{c})\right) = 0.5(1 - 2\ IBS + 1) = 1 - IBS. \text{ (Eq. 6)}$$

Note that this equivalence is conserved whether we consider ante- or post-selection parental contributions ($\boldsymbol{c}$), respectively in OCS or in UCPC based OCS.
