## Supplementary material for "Improving short and long term genetic gain by accounting for within family variance in optimal cross selection": File S3

File S3 - Supporting R code

*Allier Antoine, Christina Lehermeier, Alain Charcosset, Laurence Moreau and Simon Teyssèdre*

This document illustrates how the UCPC (Usefulness Criterion Parental Contribution, described in Allier et al. (2019) in G3 5:1469-1479, doi: https://doi.org/10.1534/g3.119.400129) is used to predict the expected gain and parental contributions after progeny selection for a cross between two homozygous parental lines. We further extend it to the evaluation of a set of crosses (**nc**) underlying the UCPC based optimal cross selection.

### 1 Load examplary data

For this documentation, we load a simulated maize data set from the `synbreedData` R package, which includes genotypic data and a genetic map.

```
rm(list=ls())
library(synbreed)
library(synbreedData)
set.seed(1993)
# Use simulated maize data set from synbreedData package as example data
data(maize)
# Convert genotypes into -1, 1 coding
geno <- maize$geno*2-1
# Set a genetic map object
map <- data.frame(CHROMOSOME=maize$map$chr,
                  POSITION=maize$map$pos,
                  MARKER=rownames(maize$map))
```

### 2 Usefulness criterion and parental contribution (UCPC) for a single two-way cross

We sample two homozygous parents further referred to as $P_1$ and $P_2$.

```
# Sample illustrative parental lines
xP1 <- geno[10,]
xP2 <- geno[200,]
```

#### 2.1 Genotypic covariance in doubled haploid progeny (DH-1) of $P_1$ x $P_2$

The following function computes the genotypic covariance matrix $\mathbf{\Sigma}$ in doubled haploid (DH) progeny derived from the cross between homozygous lines $P_1$ and $P_2$ as described in Lehermeier et al. (2017) Genetics 207:1651-1661.

Function arguments:

- ObjectGenoP1: Line vector of genotypes at loci of parent $P_1$ (coded -1/1)
- ObjectGenoP2: Line vector of genotypes at loci of parent $P_2$ (coded -1/1)
- ObjectMap: Data.frame of the genetic map with columns: CHROMOSOME, POSITION and MARKER

Value:

- Returns the covariance matrix of genotypes in DH progeny of the cross $P_1$ x $P_2$

```
GenCovProgeny   <- function(ObjectGenoP1, ObjectGenoP2, ObjectMap){
  # Compute expected frequency of recombinants c1
  myDist <- sapply(1:nrow(ObjectMap),
```

```
                    function(x) abs(ObjectMap$POSITION[x] - ObjectMap$POSITION))
  myCHR1 <- do.call(rbind, lapply(1:nrow(ObjectMap),
                                   function(x) rep(ObjectMap$CHROMOSOME[x],nrow(ObjectMap))))
  myCHR2 <- t(myCHR1)
  c1 <- 0.5*(1-exp(-2*(myDist/100)))
  c1[myCHR1!=myCHR2] <- 0.5
  Recomb <- (1-2*c1)
  # Compute disequilibrium
  tmp <- rbind(ObjectGenoP1,ObjectGenoP2)/2
  D <- crossprod(scale(tmp,scale=F))/2
  # Sigma matrix
  MyVarCov <- 4*D*Recomb
  return(MyVarCov)
}
```

The genotypic covariance matrix in the illustrative example is computed as:

```
Sigma <- GenCovProgeny(ObjectGenoP1 = xP1,
                       ObjectGenoP2 = xP2,
                       ObjectMap = map)
```

#### 2.2 Ante-selection: progeny means and co-variances

##### 2.2.1 Definition of marker effects

Let us simulate the column vector of effects $\beta_T$ for the performance trait $T$ as:

```
# Simulate marker effects (can be replaced by estimated marker effects)
BetaT <- matrix(rnorm(ncol(geno),0,sqrt(0.05)), ncol=1)
```

It results in the parent $P_1$ and $P_2$ breeding values:

```
# P1 performance is:
round(xP1%*%BetaT, digits=3)[1,1]
```

```
## [1] -8.438
```

```
# P2 performance is:
round(xP2%*%BetaT, digits=3)[1,1]
```

```
## [1] -8.836
```

The following function defines the column vector of marker effects $\beta_{C1}$ to follow $P_1$ identity by state (IBS) contribution to progeny considering only polymorphic loci between parents $P_1$ and $P_2$.

Function arguments:

- ObjectGenoP1: Line vector of genotypes at loci of parent $P_1$ (coded -1/1)
- ObjectGenoP2: Line vector of genotypes at loci of parent $P_2$ (coded -1/1)

Value:

- Returns a column vector $\beta_{C1}$ of effects to follow $P_1$ parental IBS contribution to progeny considering only polymorphic loci between $P_1$ and $P_2$.

```
GetBetaC1 = function(ObjectGenoP1, ObjectGenoP2){
  X1tmp <- matrix(ObjectGenoP1, ncol=1)
  X2tmp <- matrix(ObjectGenoP2, ncol=1)
```

```
  tmp <- X1tmp-X2tmp
  return(tmp/crossprod(tmp)[1])
}
BetaC1 <- GetBetaC1(ObjectGenoP1 = xP1,
                    ObjectGenoP2 = xP2)
```

It results in the parent $P_1$ contribution to parents:

```
# P1 contribution to itself is:
round(xP1%*%BetaC1 + 0.5, digits=3)[1,1]
```

```
## [1] 1
```

```
# P1 contribution to P2 is:
round(xP2%*%BetaC1 + 0.5, digits=3)[1,1]
```

```
## [1] 0
```

##### 2.2.2 Progeny means

The effects $\beta_T$ and $\beta_{C1}$ are used to compute progeny means ($\mu_T$ and $\mu_{C1}$) before selection:

```
# Progeny mean performance before selection is:
MuT <- 0.5*(xP1%*%BetaT+xP2%*%BetaT)
round(MuT, digits=3)[1,1]
```

```
## [1] -8.637
```

```
# Progeny mean P1 contribution before selection is:
MuC1 <- 0.5*(xP1%*%BetaC1 + xP2%*%BetaC1 + 1)
round(MuC1, digits=3)[1,1]
```

```
## [1] 0.5
```

##### 2.2.3 Progeny co-variances

The following function computes the genetic co-variances of two traits in progeny based on marker effects and the genotypic covariance matrix in progeny $\mathbf{\Sigma}$.

Function arguments:

- ObjectBeta1: Column vector of marker effects for trait 1 ($\beta_1$)
- ObjectBeta2: Column vector of marker effects for trait 2 ($\beta_2$)
- ObjectSigma: Covariance matrix of genotypes in progeny ($\mathbf{\Sigma}$)

Value:

- Returns the genetic covariance $\beta_1'\mathbf{\Sigma}\beta_2 = \beta_2'\mathbf{\Sigma}\beta_1$.

```
VarCovProgeny = function(ObjectBeta1, ObjectBeta2, ObjectSigma){
  crossprod(ObjectBeta1,ObjectSigma%*%ObjectBeta2)
}
```

The genetic variance in progeny for the performance trait $T$ ($\sigma_T^2$) is:

```
VarT <- VarCovProgeny(ObjectBeta1 = BetaT,
                      ObjectBeta2 = BetaT,
                      ObjectSigma = Sigma)
round(VarT, digits=3)[1,1]
```

```
## [1] 16.94
```

The genetic variance in progeny for $P_1$ contribution trait ($\sigma^2_{C1}$) is:

```
VarC1 <- VarCovProgeny(ObjectBeta1 = BetaC1,
                       ObjectBeta2 = BetaC1,
                       ObjectSigma = Sigma)
round(VarC1, digits=3)[1,1]
```

```
## [1] 0.014
```

The genetic covariance between performance trait and $P_1$ contribution trait ($\sigma_{T,C1}$) is:

```
CovTC1 <- VarCovProgeny(ObjectBeta1 = BetaT,
                        ObjectBeta2 = BetaC1,
                        ObjectSigma = Sigma)

round(CovTC1, digits=3)[1,1]
```

```
## [1] 0.048
```

##### 2.3 Post-selection: usefulness criterion and parental contribution (UCPC)

The following function computes the usefulness criterion parental contribution (UCPC) for a single cross $P_1$ x $P_2$ based on progeny means and genetic co-variances previously predicted and a within cross selection of the *pSel* most performant progeny.

Function arguments:

- ObjectMuT: Progeny mean for the performance trait $T$ ($\mu_T$)
- ObjectVarT: Progeny genetic variance for the performance trait $T$ ($\sigma^2_T$)
- ObjectMuC1: Progeny mean for the $P_1$ contribution (i.e. $\mu_{C1} = 0.5$ for two-way cross)
- ObjectCovTC1: Progeny genetic covariance between the performance and the $P_1$ contribution ($\sigma_{T,C1}$)
- ObjectpSel: Percentage of selected progeny within the family
- h: Selection accuracy (by default h=1)

Value:

- Returns a data.frame giving the progeny mean performance before selection ($\mu_T$), the usefulness criterion for the performance trait $T$ ($UC^{(i)}_T$), the expected $P_1$ and $P_2$ contributions to progeny before selection ($c_1$, $c_2$) and the expected $P_1$ and $P_2$ contributions to the selected fraction of progeny ($c^{(i)}_1$, $c^{(i)}_2$).

```
GetUCPC = function(ObjectMuT, ObjectVarT, ObjectMuC1, ObjectCovTC1,
                   ObjectpSel, h=1){
  i <- dnorm(qnorm(1-ObjectpSel))/ObjectpSel
  # Usefulness criterion on T
  UCT <- ObjectMuT+i*h*sqrt(ObjectVarT)
  # Correlated response to selection for C1
  C1sel <- ObjectMuC1+i*h*ObjectCovTC1/sqrt(ObjectVarT)
  return(data.frame(MuT = ObjectMuT,
                    UCT = UCT,
                    MuC1 = ObjectMuC1,
                    MuC2 = 1-ObjectMuC1,
                    SelC1 = C1sel,
                    SelC2 = 1-C1sel))
  }

UCPC <- GetUCPC(ObjectMuT = MuT, ObjectVarT = VarT, ObjectMuC1 = MuC1,
```

```
                ObjectCovTC1 = CovTC1, ObjectpSel = 0.05)

UCPC
```

```
##          MuT        UCT MuC1 MuC2    SelC1    SelC2
## 1 -8.637359 -0.1475092  0.5  0.5 0.524217 0.475783
```

### 3 UCPC to evaluate the interest of a set of two-way crosses

In optimal cross selection, we evaluate a set of crosses as a whole instead of independent single crosses.

```
# Sample of a set of twenty two-way crosses
SampledP1 <- sample(1:nrow(geno),20, replace = TRUE)
SampledP2 <- sample(setdiff(1:nrow(geno),SampledP1),20, replace = TRUE)
# Create a cross object: with parents and within cross percentage of selected progeny
Crosses = data.frame(PARENT1 = SampledP1,
                     PARENT2 = SampledP2,
                     PSelect = 0.05,
                     stringsAsFactors = FALSE)
```

#### 3.1 Compute the UCPC for several two-way crosses

The following function implements the UCPC previously defined for a single cross in a loop for several crosses.

Function arguments:

- ObjectCrosses: Data.frame of crosses with columns: PARENT1, PARENT2 and PSelect giving respectively the first and second parent of the cross and the within family selected fraction of progeny
- ObjectBetaT: Column vector of trait performance marker effects $\beta_T$
- ObjectGeno: Genotype of all candidate parents in lines and markers in column (coded in -1/1)
- ObjectMap: Data.frame of the genetic map with columns: CHROMOSOME, POSITION and MARKER

Value:

- Returns a data.frame giving for every cross: the progeny mean performance before selection ($\mu_T$), the usefulness criterion for the performance trait $T$ ($UC_T^{(i)}$), the expected $P_1$ and $P_2$ contributions to progeny before selection ($c_1$, $c_2$) and the expected $P_1$ and $P_2$ contributions to the selected fraction of progeny ($c_1^{(i)}$, $c_2^{(i)}$).

```
GetSetUCPC = function(ObjectCrosses, ObjectBetaT, ObjectGeno, ObjectMap){
  return(do.call(rbind,lapply(1:nrow(ObjectCrosses), function(nCross){
    # Parents and within cross selection parameters
    P1 <- ObjectCrosses$PARENT1[nCross]
    P2 <- ObjectCrosses$PARENT2[nCross]
    pSel <- ObjectCrosses$PSelect[nCross]
    # Genotype of parents
    xP1 <- ObjectGeno[P1,]
    xP2 <- ObjectGeno[P2,]
    # Get Sigma matrix
    Sigma <- GenCovProgeny(ObjectGenoP1 = xP1,
                           ObjectGenoP2 = xP2,
                           ObjectMap = ObjectMap)
    # Construct C1 effects
    BetaC1 <- GetBetaC1(ObjectGenoP1 = xP1,
```

```
                        ObjectGenoP2 = xP2)
    # Get ante-selection means and co-variances
    MuT <- 0.5*(xP1%*%ObjectBetaT+xP2%*%ObjectBetaT)
    VarT <- VarCovProgeny(ObjectBeta1 = ObjectBetaT,
                          ObjectBeta2 = ObjectBetaT,
                          ObjectSigma = Sigma)
    MuC1 <- 0.5*(xP1%*%BetaC1 + xP2%*%BetaC1 + 1)
    CovTC1 <- VarCovProgeny(ObjectBeta1 = ObjectBetaT,
                            ObjectBeta2 = BetaC1,
                            ObjectSigma = Sigma)
    # Get the UCPC considering within family selection
    UCPC <- GetUCPC(ObjectMuT = MuT, ObjectVarT = VarT,
                    ObjectMuC1 = MuC1, ObjectCovTC1 = CovTC1,
                    ObjectpSel = pSel)

    return(cbind(data.frame(Cross = paste0(P1,"x",P2),
                            PARENT1 = P1,
                            PARENT2 = P2,
                            PSel = pSel,
                            stringsAsFactors = FALSE),
                 UCPC))
  }))
}

SetUCPC <- GetSetUCPC(ObjectCrosses = Crosses, ObjectGeno = geno,
                      ObjectBetaT = BetaT, ObjectMap = map)
SetUCPC[1:5,-c(2,3)]
```

```
##       Cross PSel       MuT       UCT MuC1 MuC2     SelC1     SelC2
## 1   20x315 0.05 -9.797972 -1.656583  0.5  0.5 0.5473419 0.4526581
## 2 702x1227 0.05  1.580834 10.411533  0.5  0.5 0.4579492 0.5420508
## 3  822x367 0.05 -6.345915  2.739877  0.5  0.5 0.4696598 0.5303402
## 4  186x182 0.05 -4.039036  2.027207  0.5  0.5 0.5029892 0.4970108
## 5  626x913 0.05  2.735078 14.270303  0.5  0.5 0.4924262 0.5075738
```

#### 3.2 Evaluate a set of crosses for expected genetic gain and genetic diversity

To evaluate this set of crosses we need to compute the gain term $V^{(i)}(\mathbf{nc})$ and diversity constraint term $D^{(i)}(\mathbf{nc})$ depending on the within family selection intensity $i$.

The IBS coancestry among candidate parents is defined using the following function.

Function arguments:

- ObjectGeno: Genotype of all candidate parents in lines and markers in column (coded in -1/1)

Value:

- Returns a matrix of IBS coancestry among candidate parents.

```
GetIBS = function(ObjectGeno){
  0.5*(tcrossprod(ObjectGeno)/ncol(ObjectGeno)+1)
}
```

Then, the following function is used to evaluate the set of crosses either accounting (UCPC) or not (OCS) for within family selection.

Function arguments:

- ObjectUCPC: UCPC data.frame obtained previously for the set of crosses
- ObjectGeno: Genotype of all candidate parents in lines and markers in column (coded in -1/1)

Value:

- Returns a data.frame with the number of crosses in the set ("nCROSS"), the number of unique parents ("nUniqParents"), $V^{(i=0)}(\mathbf{nc})$ and $D^{(i=0)}(\mathbf{nc})$ not accounting for within family selection ("OCS_Vnc" and "OCS_Dnc"), $V^{(i)}(\mathbf{nc})$ and $D^{(i)}(\mathbf{nc})$ accounting for within family selection ("UCPC_Vnc" and "UCPC_Dnc").

```
EvaluateSetUCPC = function(ObjectUCPC, ObjectGeno){
  # Incidence matrices Z1 and Z2
  UniqueParents <- unique(c(ObjectUCPC$PARENT1,ObjectUCPC$PARENT2))
  Z1 <- matrix(0,ncol=nrow(ObjectUCPC),nrow=length(UniqueParents))
  Z2 <- matrix(0,ncol=nrow(ObjectUCPC),nrow=length(UniqueParents))
  invisible(lapply(1:length(UniqueParents), function(x){
    Z1[x,ObjectUCPC$PARENT1==UniqueParents[x]] <<-1
    Z2[x,ObjectUCPC$PARENT2==UniqueParents[x]] <<-1
  }))
  # Compute IBS matrix
  K <- GetIBS(ObjectGeno = ObjectGeno[UniqueParents,])
  # UCPC based OCS: D(nc) and V(nc) terms accounting for within family selection
  c1 <- matrix(ObjectUCPC$SelC1, ncol=1)
  c2 <- matrix(ObjectUCPC$SelC2, ncol=1)
  c <- (Z1%*%c1+Z2%*%c2)/nrow(ObjectUCPC)
  UCPC_Dnc <- 1-crossprod(c,K%*%c)
  UCPC_Vnc <- mean(ObjectUCPC$UCT)
  # classical OCS: D(nc) and V(nc) terms not accounting for within family selection
  c1 <- matrix(ObjectUCPC$MuC1, ncol=1)
  c2 <- matrix(ObjectUCPC$MuC2, ncol=1)
  c <- (Z1%*%c1+Z2%*%c2)/nrow(ObjectUCPC)
  OCS_Dnc <- 1-crossprod(c,K%*%c)
  OCS_Vnc <- mean(ObjectUCPC$MuT)

  return(data.frame(nCROSS = nrow(ObjectUCPC),
                    nUniqParents = length(UniqueParents),
                    OCS_Vnc = OCS_Vnc,
                    OCS_Dnc = OCS_Dnc,
                    UCPC_Vnc = UCPC_Vnc,
                    UCPC_Dnc = UCPC_Dnc,
                    stringsAsFactors = FALSE))
}
```

```
SetEval <- EvaluateSetUCPC(ObjectUCPC = SetUCPC, ObjectGeno = geno)
SetEval
```

```
##   nCROSS nUniqParents    OCS_Vnc   OCS_Dnc UCPC_Vnc  UCPC_Dnc
## 1     20           40 -0.8399918 0.3373086 8.155683 0.3367695
```
